# Ligand-induced segregation from large cell-surface phosphatases is a critical step in γδ TCR triggering

**DOI:** 10.1101/2023.08.23.554524

**Authors:** Fenglei Li, Sobhan Roy, Jacob Niculcea, Keith Gould, Erin J. Adams, P. Anton van der Merwe, Kaushik Choudhuri

**Affiliations:** Department of Microbiology and Immunology, University of Michigan Medical School, Ann Arbor, MI 48109, USA; Department of Biochemistry and Molecular Biology, University of Chicago, Chicago, IL 60637, USA; Department of Infectious Diseases, Imperial College London, London, UK; Sir William Dunn School of Pathology, University of Oxford, Oxford, UK

## Abstract

Gamma/delta (γδ) T cells are unconventional adaptive lymphocytes that recognize structurally diverse ligands *via* somatically-recombined antigen receptors (γδ TCRs). The molecular mechanism by which ligand recognition initiates γδ TCR signaling, a process known as TCR triggering, remains elusive. Unlike αβ TCRs, γδ TCRs are not mechanosensitive, and do not require coreceptors or typical binding-induced conformational changes for triggering. Here, we show that γδ TCR triggering by nonclassical MHC class Ib antigens, a major class of ligands recognized by γδ T cells, requires steric segregation of the large cell-surface phosphatases CD45 and CD148 from engaged TCRs at synaptic close contact zones. Increasing access of these inhibitory phosphatases to sites of TCR engagement, by elongating MHC class Ib ligands or truncating CD45/148 ectodomains, abrogates TCR triggering and T cell activation. Our results identify a critical step in γδ TCR triggering and provide insight into the core triggering mechanism of endogenous and synthetic tyrosine-phosphorylated immunoreceptors.

## Introduction

γδ T cells are unconventional lymphocytes with innate-like characteristics that play key roles in anti-microbial immunity and inflammation^1, 2, 3^. They are activated by structurally diverse ligands *via* distinct somatically-recombined γδ T cell antigen receptors (γδ TCR)^4^. While many of the ligands that activate γδ T cells remain poorly defined, MHC class Ib molecules constitute a major class of membrane-tethered ligands that are recognized by both murine and human γδ T cells^5^. These minimally polymorphic nonclassical MHC class I antigens are widely expressed on immune and non-immune cells, and are upregulated on cell surfaces by intracellular stress, inflammatory mediators, or exposure to microbial components. The γδ TCR is a multi-subunit receptor complex composed of an antigen-binding γδ heterodimer assembled with dimeric CD3 and TCR-ζ signaling subunits^6, 7^ that contain immunoreceptor tyrosine-based activation motifs (ITAMs)^8^ in their cytoplasmic domains. Engagement of the TCR by cognate ligands initiates intracellular signaling by inducing phosphorylation of TCR ITAMs, a process known as TCR triggering^9^. The mechanism of γδ TCR triggering is not known, although several mechanisms implicated in αβ TCR triggering are likely not involved. A characteristic binding-induced conformational change in the cytoplasmic tail of CD3ε subunits of αβ TCRs^10^, which recruits the adaptor protein Nck, does not occur in the murine G8 γδ TCR following engagement by its cognate ligand T22^11^ – a murine MHC class Ib antigen. Moreover, the homologous segment of the mechanosensitive TCR Cβ F-G loop of αβ TCRs is truncated in TCR Cγ, suggesting that γδ TCRs may not be mechanically activated^12^. This was recently confirmed by direct force measurements of the human γδ TCR DP10.7, interacting with its cognate ligand, the sulfoglycolipid sulfatide complexed with the human MHC class Ib molecule CD1d (sulfatide-CD1d)^13^. Despite effective triggering, TCR DP10.7 binds only transiently to sulfatide-CD1d under load, and lacks the characteristic catch bond force profile that correlates with pMHC-mediated αβ TCR triggering. Mechanisms of triggering that rely on coreceptor-mediated TCR dimerization/aggregation are also unlikely to operate in γδ T cells, as they exit the thymus at the CD4/CD8 double-negative stage during thymic development, and are predominantly coreceptor negative^14^.

Here, we investigate whether γδ TCR triggering is instead more reliant on the kinetic-segregation (K-S) model of TCR triggering^15^, which does not require conformational changes, mechanical forces or coreceptor binding. The K-S model proposes that, due to the relatively small size of TCRs and their membrane-tethered ligands, TCR engagement takes place within zones of close contact between T cells and antigen-presenting cells (∼15 nm), from which T cell surface phosphatases with large ectodomains, such as CD45 and CD148, are sterically excluded. The resulting shift in local kinase-phosphatase balance allows stable phosphorylation of engaged TCRs by the Src-family kinase Lck^16^, thus initiating signaling. We show that increasing access of CD45 and CD148 to sites of γδ TCR engagement abrogates triggering and T cell activation, in keeping with the K-S model.

## Results

### Coreceptors and CD3ε conformational change do not contribute to γδ TCR triggering

In order to establish which TCR triggering mechanism(s) operate in γδ T cells, we chose to focus initially on the well-characterized murine G8 γδ TCR and its ligand T22 as a model experimental system^17^. We engineered an expression construct comprising a single-chain dimer (SCD) of T22 in which β2M was fused to the N-terminus of the T22 α-chain via a 10-residue glycine/serine linker (Fig. 1a, Extended Data Fig. 1a). The T22 SCD construct was transfected into CHO cells, and stable transfectants were sorted by FACS, using T22-specific hamster mAb 7H9 (Gift from Y.H. Chien, Stanford) and a fluorescently labelled secondary antibody, to obtain sorted CHO cells with well-defined cell-surface levels of T22 (T22-CHO) (Fig. 1b). T cell hybridomas expressing cognate G8 γδ TCRs (G8 T cells), or another T22-specific TCR – KN6^18^, were robustly activated by co-culture with T22-CHO cells as measured by IL-2 release in culture supernatants (Fig. 1c, e). Neither G8 nor KN6 T cell hybridomas expressed detectable levels of CD8 or CD4 on their surface, precluding a role for coreceptors in their activation (Extended Data Fig. 1b,c).

**Figure 1.**
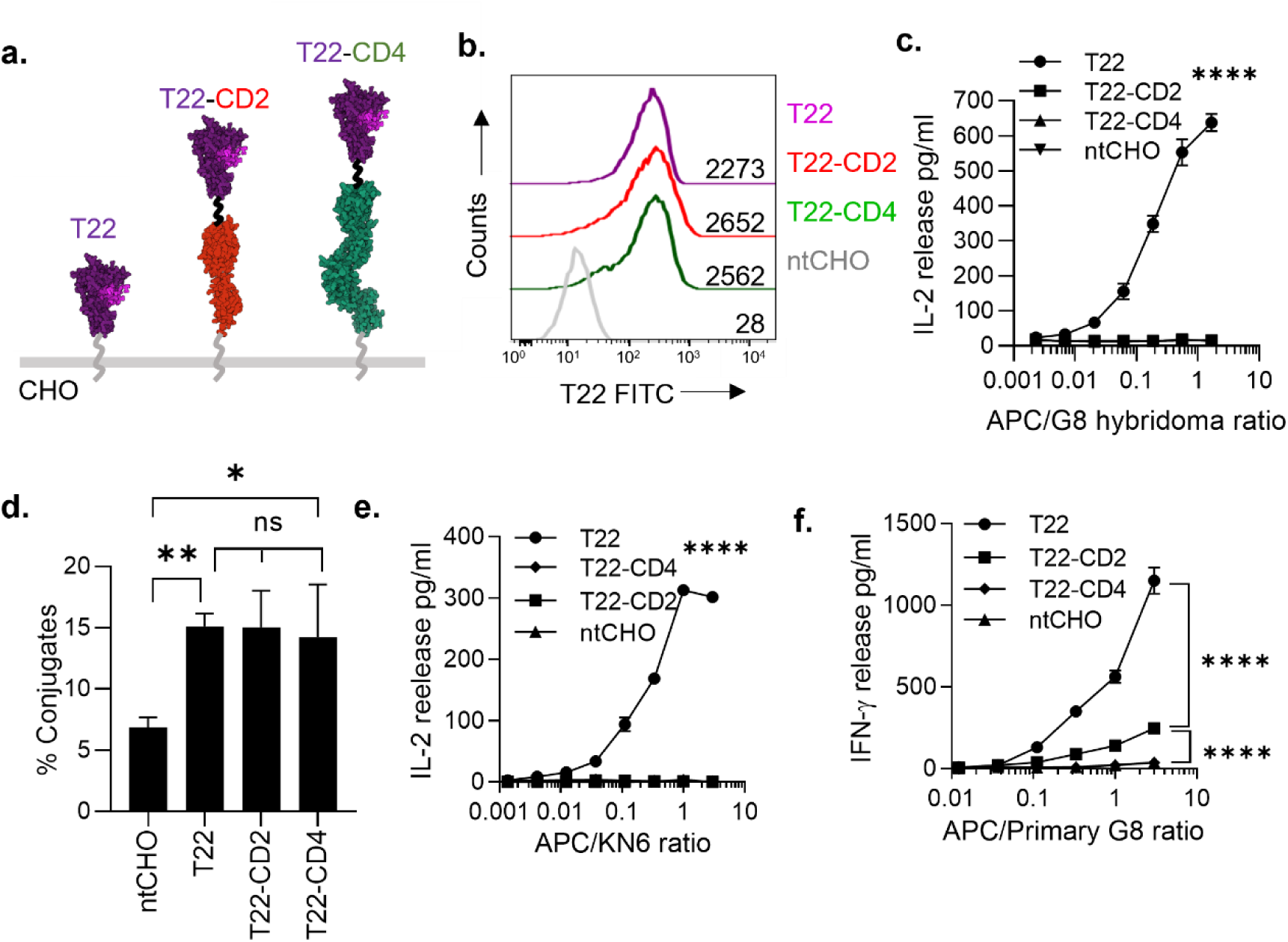
Elongation of murine MHC class Ib molecule T22 abrogates cognate γδ T cell activation. **a**. Structure-based schematic of single-chain constructs of T22 and elongated forms of T22. β2M (pink) is attached to the N-terminus of T22 using a glycine/serine linker. In elongated forms of T22, spacers derived from human CD2 or CD4 are joined to the C-terminus of T22 by glycine/serine linkers (black). All constructs are tethered to the plasma membrane by the mouse H-2D^b^ stalk segment (grey). **b.** Surface expression of the indicated T22 constructs in stably transfected CHO cells or untransfected CHO cells (ntCHO), detected using a T22-specific antibody and fluorescently labeled secondary anti-hamster antibody. **c.** IL-2 release by cognate γδ G8 T cell hybridomas (10^4^/well), co-cultured with increasing numbers of CHO cells expressing comparable levels of the indicated T22 constructs, or with untransfected CHO cells (nt). Results are representative of 3 independent experiments. **d.** Cell-cell adhesion assay using a 1:1 mixture of fluorescently-labeled G8 T cell hybridomas (CMFDA – green fluorescence) and CHO cells (Cell Tracker Deep Red – red fluorescence) expressing the indicated T22 constructs. Heteroconjugates were detected by flow-cytometry as percentage of events ‘double-positive’ for both G8 and CHO cell-associated fluorescence. **e.** IL-2 release from KN6 γδ T cell hybridomas in response to increasing numbers CHO cells expressing the indicated T22 constructs. **f.** IFN-γ release by primary T22-specific primary γδ T cells isolated from *Rag^-^/^-^ BALB/c* mice transgenic for the G8 γδ TCR. Data points in ***c.-f.*** are means ± SD. ****, P < 0.0001; **, P < 0.01; *, P < 0.05; ns, not significant (P > 0.05); P values are corrected for all-pairwise comparisons. Panels are representative results of 4 independent experiments unless otherwise indicated.

The Nck-recruiting CD3ε conformational change does not occur in G8 TCRs following engagement by cell-expressed T22^11^, although it is unclear whether this is true more generally for γδ TCRs. We therefore tested for this conformational change in a second, lower affinity (K_D_ 16 μM)^19^, T22-specific murine γδ TCR – KN6^18^. KN6 T cell hybridomas were brought into contact with T22-expressing CHO cells at 4°C to induce intercellular contact, and activated by incubation at 37°C for 10 minutes. Cells lysates were probed using a well-established TCR pulldown assay^10^, modified to detect conformation-sensitive TCR binding to biotinylated Nck SH3.1 peptides immobilised on streptavidin-coated ferromagnetic beads. As with the G8 TCR^11^, KN6-associated TCRζ was not immunoprecipitated by bead-immobilised Nck3.1 peptides following engagement by T22-CHO cells, or by soluble T22 tetramers (Extended Data Fig. 2a), but was readily detected when KN6 hybridomas were artificially stimulated with activating anti-CD3ε mAb 2C11 (Extended Data Fig. 2a).

We also tested for the CD3ε conformational change in more physiological primary γδ T cells, isolated from G8 TCR transgenic mice bred on a *Balb/c Rag^-/-^* background^20, 21, 22^ (Gift from YH Chien, Stanford). Primary G8 γδ T cells were isolated from spleen and lymph nodes and enriched by negative selection to >95% purity, as measured by labelling with a fluorescent T22-tetramer (PE), and detected using flow-cytometry (Extended Data Fig. 2b). Unlike αβ T cells, saturating concentrations of soluble T22 tetramers negligibly activated G8 T cells as measured by IL-2 release (Extended Data Fig. 2d), although they were readily activated by T22 monomers adsorbed to flat-bottomed culture wells (Extended Data Fig. 2e), suggesting that G8 T cell activation of was contact-dependent. As with G8 T cell hybridomas, primary G8 T cells were robustly activated by T22-CHO cells, as measured by IFNγ release (Fig. 1f). However, the CD3ε conformational change was not detected, either by Nck SH3.1-mediated TCR pulldown (Extended Data Fig. 2c), or by labelling with the conformation-sensitive anti-CD3ε mAb APA1/1^23^ (Extended Data Fig. f,g). Taken together, these results show that typical ligand-induced TCR conformational changes are absent in murine γδ T cells.

### T22 ectodomain size but not flexibility critically affects cognate γδ T cell activation

We next asked whether TCR triggering proceeded according to kinetic-segregation (K-S) model in γδ T cells. To test this, we adapted a powerful approach employed in our previous studies of the K-S model in αβ TCR triggering^24, 25^, in which we disrupted close contacts at T cell synapses using elongated cognate pMHC ligands containing rigid immunoglobulin superfamily (IgSF) domain spacers in the membrane-proximal stalk region. We used this approach to engineer chimeric expression constructs of elongated versions of T22 SCD by inserting the ectodomain sequence of human CD2 (2 IgSF domains) or human CD4 (4 IgSF domains) between the C-terminus of the T22 ectodomain and the N-terminus of the H-2D^b^ stalk and transmembrane segment (Fig. 1a). Stable CHO cell transfectants expressing T22 and elongated forms of T22 were sorted for comparable surface expression and co-cultured with G8 T cell hybridomas (Fig. 1b). Robust and titratable IL-2 release from G8 T cell hybridomas was observed in response to T22-expressing CHO cells (Fig. 1c). However, IL-2 release was completely abrogated by T22 elongation with CD2 or CD4 spacers (Fig. 1c). This was not due to impaired TCR binding, as G8 T cell adhesion to CHO cells expressing T22 constructs was unaffected by T22 elongation, as measured by a flow cytometry-based adhesion assay (Fig. 1d)^24^. Taken together, these results show that TCR-mediated activation of γδ T cells, as measured by ‘downstream’ readouts of TCR signaling, is exquisitely sensitive to ligand elongation.

It remained possible that, despite effective binding, the short 5 residue glycine/serine interdomain peptide linker (g/s5) connecting the IgSF spacers to T22 may have constrained it in orientations that inhibited TCR triggering, We therefore substituted the 5g/s linker with a more flexible 10 residue glycine/serine linker (T22-g/s10) (Extended Data Fig. 3a,b)^26, 27^. We also added a g/s10 linker to the stalk of unelongated T22 (T22-g/s10), to test whether increasing native ligand flexibility affected G8 T cell activation (Extended Data Fig. 3a,b). Introduction of the g/s10 linker into native-sized T22 did not affect adhesion (Extended Data Fig. 3c) or activation (Extended Data Fig. 3d) of G8 T cells hybridomas. Conversely, replacement of the short interdomain linker connecting T22 to the CD4 spacer with the g/s10 linker failed to rescue G8 T cell activation (Extended Data Fig. 3d), demonstrating that increased ectodomain flexibility of T22 constructs did not affect their activity or intermembrane spacing at synaptic contacts.

### Graded attenuation of γδ TCR triggering by progressive T22-elongation

To investigate whether the observed reduction in γδ T cell activation in response to T22 elongation was due to reduced TCR triggering, we directly assayed T22-induced tyrosine phosphorylation of the TCRζ chain^8, 24^. T cells were centrifuged at 4°C with CHO cells expressing T22 SCD, or untransfected CHO cells as a control, to induce intercellular contact, and incubated at 37°C for 2-10 minutes to initiate signaling. TCRζ was immunoprecipitated from sample lysates and tyrosine phosphorylation detected by SDS-PAGE and immunoblotting. A robust and sustained increase in TCRζ phosphorylation was detectable in G8 T cells within 2 minutes of co-culture with T22-expressing CHO cells, which peaked at 5 minutes, and remained phosphorylated after 10 minutes of incubation, with a progressive transition from the partially phosphorylated p21 phosphoform to the fully phosphorylated p23 phosphoform (Fig. 2a)^8^. In marked contrast, CHO cells expressing T22-CD2 and T22-CD4 barely elicited detectable TCRζ phosphorylation, which was indistinguishable from baseline levels within 5 minutes of coculture (Fig. 2a). These results directly demonstrate that G8 TCR phosphorylation is critically sensitive to T22 elongation, resulting in near complete abrogation of TCR triggering.

**Figure 2.**
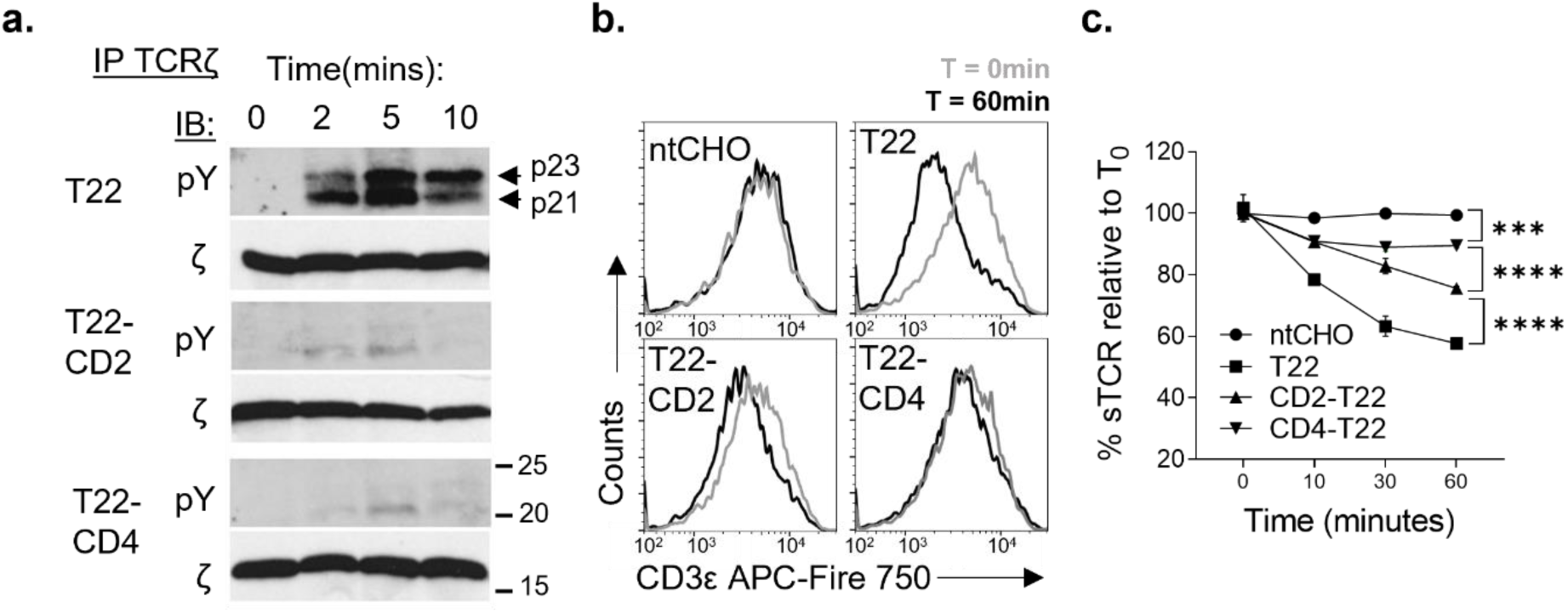
G8 γδ TCR triggering is abolished by T22 elongation. **a**. Timeseries of TCR-ζ phosphorylation, measured by immunoprecipitation of TCR-ζ, followed by SDS-PAGE and immunoblotting with an anti-phosphotyrosine antibody (pY) to detect tyrosine-phosphorylated isoforms of TCR-ζ (arrows). p21, partially phosphorylated 21 KD ζ isoform, p23, fully phosphorylated ζ isoform. Blots were reprobed to detect unphosphorylated TCR-ζ as a loading control (ζ). Results are representative of 3 independent experiments. **b.** Representative histogram plots of TCR surface downregulation in G8 γδ T cell hybridomas incubated at 1:1 ratio for 0 to 60 minutes with CHO cells expressing the indicated T22 constructs or untransfected CHO cells (ntCHO). Surface TCR was measured in unpermeabilized samples using a directly-labelled mouse CD3ε-specific antibody and detected by flow-cytometry. Overlays of 0 (grey) and 60 (black) minute timepoints are shown. **c.** Timeseries of TCR surface downregulation, in response to T22, elongated forms of T22, or untransfected CHO cells (nt). Results are presented as percentage of surface TCR fluorescence relative to T=0 levels. Results are representative of 3 independent experiments. ***, P < 0.001, ****, P < 0.0001. P values are for data points at T=60min and corrected for all pairwise comparisons.

As a second readout of TCR triggering, we measured antigen-induced surface downregulation of the TCR, which reflects the proportion of fully triggered and internalised TCR^28^. As expected, incubation of G8 T cells with T22-CHO cells resulted in clear-cut TCR downregulation, with a ∼40% reduction of surface TCR levels after 60 minutes co-culture (Fig. 2b,c). T22-CD2 and T22-CD4 elicited lower but measurable levels of TCR downregulation relative to T22, indicating weaker TCR triggering, while surface TCR levels were unchanged in T cell co-cultures with untransfected CHO cells (Fig. 2b,c). The extent of TCR downregulation decreased in a graded manner, as a function of increasing T22 ectodomain size: T22 > T22-CD2 > T22-CD4. These results show that T22 size is a critical and tunable determinant of TCR triggering.

### Increased Lck activity does not rescue G8 TCR triggering in response to elongated T22

We attempted to rescue TCR triggering in response to elongated forms of T22 by shRNA-mediated knockdown of the tyrosine phosphatase Csk, a negative regulator of Lck^29^. We achieved almost complete suppression of Csk expression in G8 T cell hybridomas (Extended Data Fig.4a), while TCR surface levels, as measured by surface labelling of CD3ε, were unaffected by shRNA expression (Extended Data Fig.4b). As expected, Csk knockdown increased phosphorylation of the positive regulatory tyrosine Y393, with reciprocal reduction in phosphorylation of the negative regulatory tyrosine Y505, indicating increased Lck activity (Extended Data Fig.4a). Surprisingly, Csk suppression had no effect on T22-mediated TCR triggering in G8 T cell hybridomas, as measured by intracellular Ca^2+^ influx following contact with T22-containing supported lipid bilayers (SLB) (Extended Data Fig. 4d). T cell activation as measured by IL-2 release was similarly unaffected by Csk knockdown (Extended Data Fig. 4c), and increased Lck activity did not rescue G8 T cell activation in response to elongated T22 ligands (Extended Data Fig. 4e). These results suggest that basal levels of Lck activity are sufficient to maximally trigger G8 TCRs. Unlike its function in tuning αβ T cell signaling^29^, Csk appears not to play a substantial role in setting TCR signaling thresholds in γδ T cells.

### T22 elongation increases intermembrane distance at γδ T cell synaptic contacts

We next investigated whether T22 elongation affected membrane positioning across synaptic contacts with G8 T cells. Using transmission electron microscopy (TEM)^24, 30^, we directly measured intermembrane distances at synaptic interfaces between G8 T cell hybridomas and CHO cells expressing T22 or T22-CD4. To achieve this, conjugates of G8 T cell hybridomas with CHO cells expressing T22 constructs were fixed after 2 minutes incubation at 37°C and processed for transmission electron microscopy (TEM). Thin-sections were screened to identify T cell:CHO cell heteroconjugates (Fig. 3a), and intermembrane distance was measured at ∼200-400 nm intervals along the contact interface (Fig. 3b,d). Intermembrane distance measurements were plotted as normalized frequency distributions of intermembrane distances with a bin of 2 nm (Fig. 3c,e). The mean intermembrane distance at interfaces between G8 T cells and T22-expressing CHO cells was 13.5 nm, consistent with the end-to-end span of the T22-G8 complex structure determined by x-ray crystallography^17^ (Fig. 3b,c). Mean intermembrane distance was significantly increased at G8 T cell interfaces with T22-CD4 expressing CHO cells to 17.5 nm, with a maximum intermembrane distance of ∼28 nm, which was consistent with the estimated dimensions of the fully-extended G8:T22-CD4 complex span (Fig. 3d,e). Substitution with a longer and more flexible g/s10 spacer into the stalk of unelongated T22, or in the interdomain linker connecting T22 and CD4 did not change the mean intermembrane spacing at synaptic contacts (Extended Data Fig. 3e), indicating that the rigid IgSF-containing spacers, and not the flexible linkers, were the primary determinants of intermembrane separation distance at synaptic contacts. These results show that apposed membranes at T22-induced intercellular contacts with G8 T cells are positioned at a distance that is consistent with the T22:G8 TCR complex span, and that T22 elongation with IgSF spacers significantly increases intermembrane distance at synaptic contacts.

**Figure 3.**
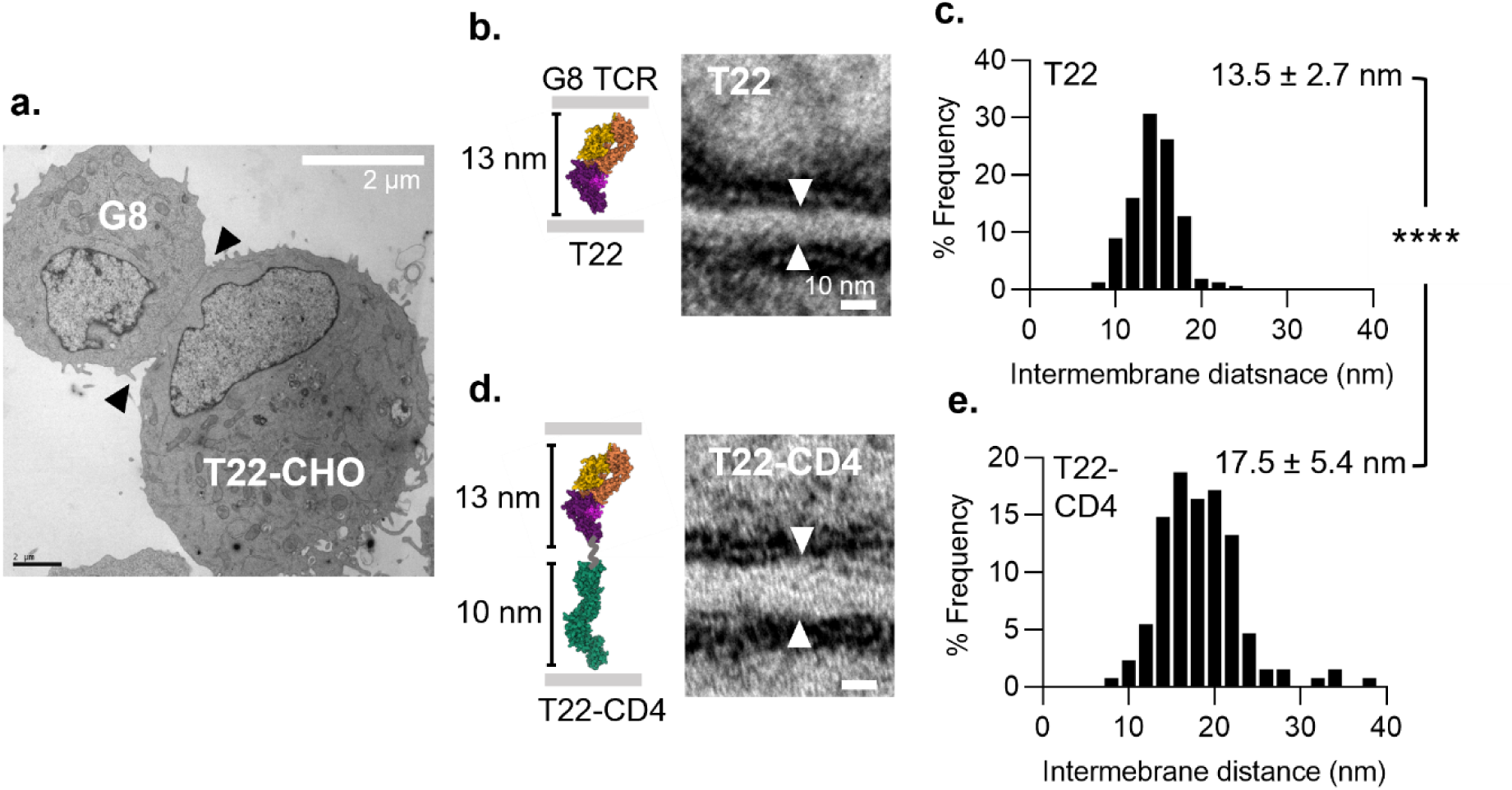
T22 elongation increases intermembrane spacing and at G8 T cell synaptic contacts. **a**. Representative heteroconjugate between a G8 T cell hybridoma and CHO cell expressing T22. Arrowheads indicate G8-CHO cell synapse along which intermembrane distance of apposed plasma membranes were measured. **b, d. Left,** Structure-based cartoon of T22 and elongated T22-CD4 constructs with estimated dimensions of ectodomains. Flexible 5 residue glycine/serine linker joining T22 to the CD4 spacer is shown in grey; **Right,** Representative high-magnification image of a region of synaptic contact between G8 T cells and CHO cells expressing T22 or T22-CD4. Arrowheads indicate region at which intermembrane distance measurements were made, at which G8 and CHO cell plasma membranes are parallel and orthogonal to the imaging plane, as indicated by their ‘tram-line’ morphology**. c, e.** Frequency distribution of intermembrane distances at synaptic contacts. A total of 189 measurements along synaptic contacts were taken from 15 conjugates between G8 T cell hybridomas and CHO cells expressing T22, and 117 measurements were taken from 18 conjugates with CHO cells expressing T22-CD4. Mean intermembrane distance ± SD is shown in each panel. ****, P < 0.0001.

### T22 ectodomain size controls γδ TCR segregation from CD45 at T cell synapses

We next sought to determine if T22 elongation affected the distribution of CD45 at G8 T cell contact interfaces. Using confocal fluorescence microscopy, we measured TCR and CD45 distributions in primary G8 T cell conjugates with CHO cells expressing T22 or T22-CD4. Conjugates were fixed after 2 minutes incubation at 37°C, and permeabilized samples were labelled with primary antibodies to TCR and CD45, followed by appropriate fluorescently-labelled secondary F(ab’)_2_ fragments, and imaged by laser-scanning confocal microscopy (Fig. 4a). Robust and comparable TCR accumulation was observed at T cell synapses with CHO cells expressing T22 and T22-CD4 (∼2-fold enrichment), indicating similar levels of TCR engagement (Fig. 4a,b). A small (∼20%) but reproducible overall depletion of CD45 was observed at synapses with T22-expressing CHO cells (Fig. 4a,c), indicating overall exclusion of CD45 from T22-mediated synaptic contacts. In contrast, CD45 levels at synapses with CHO cells expressing T22-CD4 were unchanged relative to non-synaptic plasma membrane regions, indicating increased overall access of CD45 to the T cell contact interface (Fig. 4a,c).

**Figure 4.**
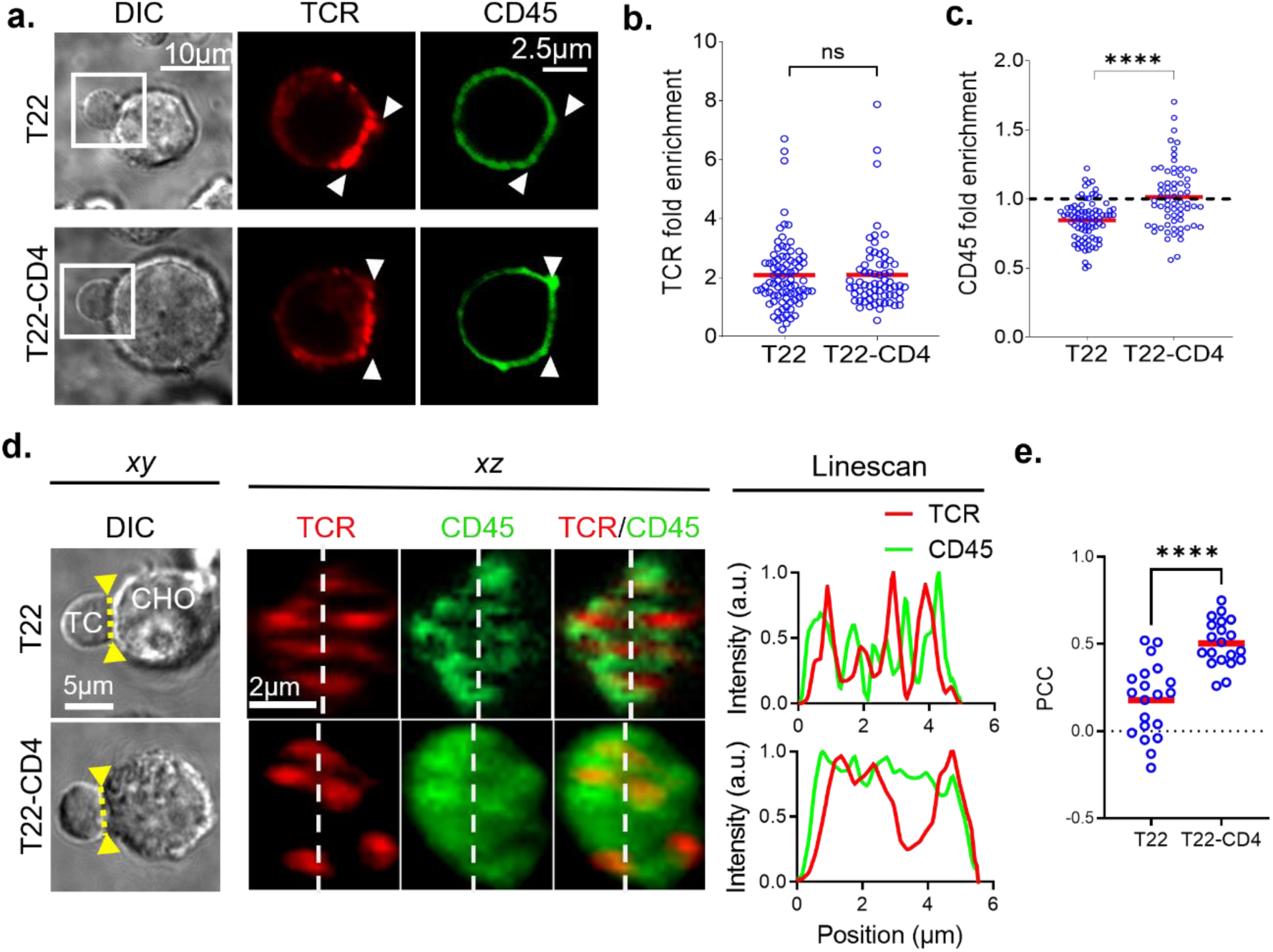
T22 ectodomain size controls CD45 segregation from TCR at G8 T cell synapses. **a**. Representative confocal fluorescence images of conjugates of primary G8 T cells and CHO cells expressing T22 or T22-CD4. TCR (red) and CD45 (green) were labeled using specific primary antibodies and fluorescently labeled secondary F(ab’)_2_ fragments. Fluorescence images of T cells in boxed regions of DIC images are shown in **right panels**. Arrowheads indicate position of synaptic interface in DIC image. **b** and **c,** Quantitation of TCR (***b.***) and CD45 (***c.***) at synaptic contacts expressed as the ratio of [mean plasma membrane fluorescence intensity at synaptic interface]/[mean plasma membrane fluorescence intensity in non-synaptic region]. Each datapoint represents measurements from one T cell:CHO cell conjugate, red bars indicate means. Results are pooled from 6 independent experiments. **d.** Representative DIC image of G8 T cell conjugates with CHO cells expressing T22 or T22-CD4 (**left**). Yellow dotted line and arrowheads indicates *xz* plane sampled in fluorescence image z-stacks of TCR (red) and CD45 (green) labeled as in ***a.*** to reconstruct *en face* views of the synaptic interface (**middle panels**). Overlays of fluorescence intensities of TCR and CD45 along linescans positioned across *en face* synapse reconstructions (white dotted line) are shown in **right panels**. **e.** Quantitation of colocalization between TCR and CD45 at reconstructed *en face* synaptic interfaces using Pearson’s correlation coefficient (PCC). ****, P < 0.0001; ns, not significant (P > 0.05); P values are corrected for all pairwise comparisons. Results are pooled from 5 independent experiments.

To measure the extent of overlap in TCR and CD45 distributions at synaptic contacts, we reconstructed *en face* (end-on) projections of synaptic interfaces from confocal z-stacks of conjugates between G8 T cells and CHO cells expressing T22 or T22-CD4^31^. Although the resolution of synapse reconstructions was limited by the axial anisotropy of the confocal point spread function, TCR microclusters, which form following TCR engagement^32^, could be resolved at contact interfaces (Fig. 4d). CD45 was segregated from TCR microclusters at contact interfaces with T22-expressing CHO cells, resulting in anticorrelation between TCR and CD45 intensities across line scans of interface regions (Fig. 4d). Characteristic ‘holes’^32^ in the CD45 distribution in regions occupied by T22-induced TCR microclusters indicated CD45 exclusion from engaged TCRs (Fig. 4d). In contrast, there was much greater overlap of CD45 distributions with TCR microclusters in *en face* reconstructions of synapses with T22-CD4-expressing CHO cells (Fig. 4d), quantitated as a highly significant increase in Pearson’s correlation coefficient (PCC), indicating greater colocalization between CD45 with TCR (Fig. 4e).

### CD45 and CD148 ectodomain size is a critical determinant of TCR triggering

While T22 elongation increases CD45 access to engaged TCRs at synaptic contacts, it remained possible that introduction of non-native IgSF spacers may also have adversely affected triggering by altering cryptic mechanical and structural properties that are important for TCR triggering. To rule this out, we examined the effects of truncating the large ectodomains of CD45 or CD148 on γδ TCR triggering by native T22. We engineered constructs of murine CD45 in which the native ectodomain was replaced with the single IgSF-containing ectodomain of rat Thy-1 (Thy1-CD45), or with the ectodomain of rat CD43 (CD43-CD45) (Fig. 5a, Extended Data Fig. 1a) which is comparable in size to CD45 (∼45 nm)^33, 34^. We also made a Thy-1-CD45 catalytically inactive mutant (Thy1-CD45*) to measure the contribution of its phosphatase activity on TCR triggering (Fig. 5a, Extended Data Fig. 1a)^35^. All constructs contained N-terminus Flag-tags for immunolabeling, cell sorting, and fluorescence imaging (Fig. 5a). CD45 expression constructs were retrovirally transduced into G8 T cell hybridomas and transductants sorted for comparable surface levels (Fig. 5b).

**Figure 5.**
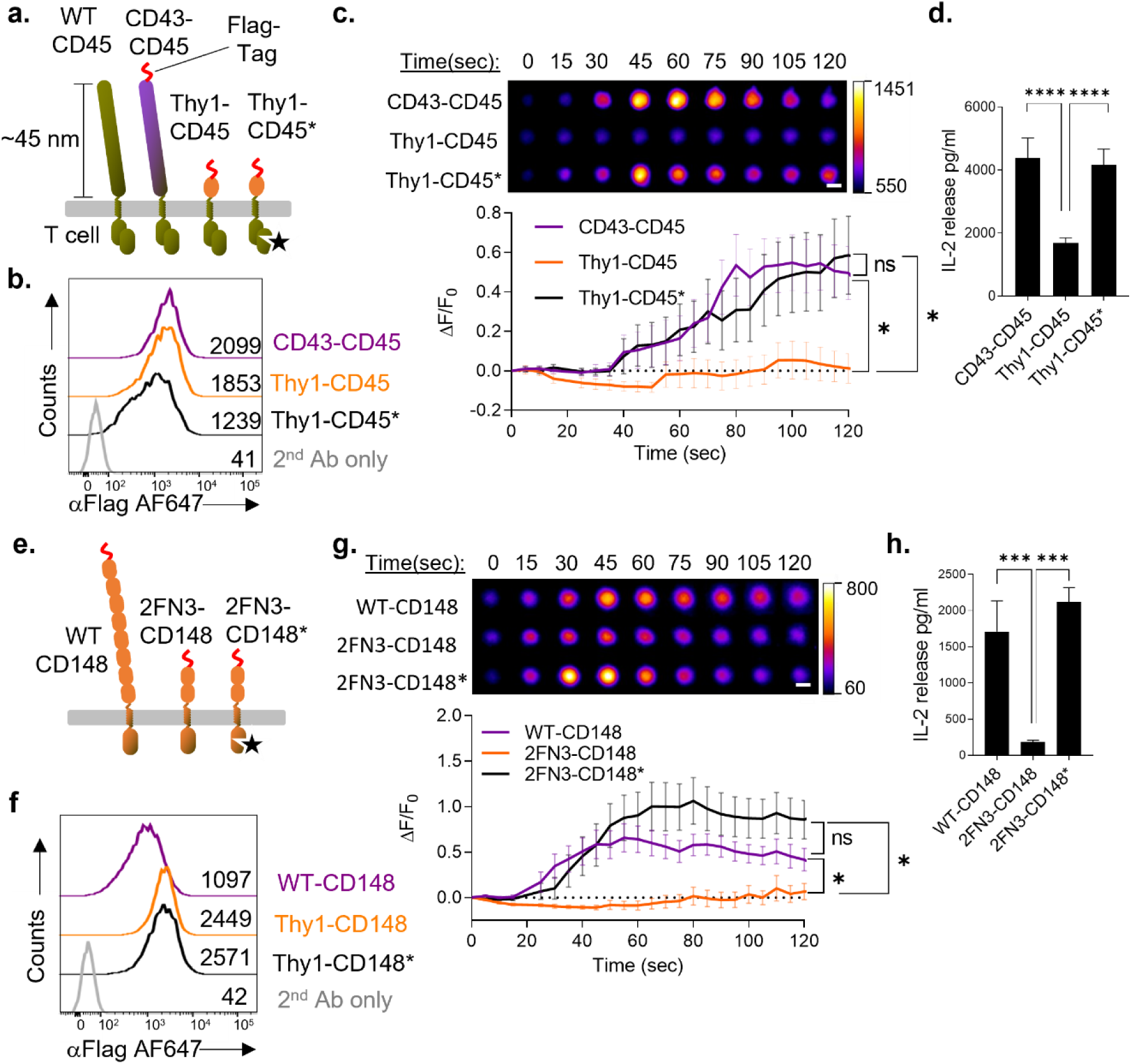
Truncation of the CD45 and CD148 ectodomains abrogates γδ TCR triggering and T cell activation. **a**. Schematic of wild-type CD45 (shown for comparison), and chimeras of the wild-type CD45 cytoplasmic domain fused to the ectodomains of CD43 (CD43-CD45) or Thy-1 (Thy1-CD45), depicted approximately to scale. Also shown is a chimera of a catalytically inactive mutant of the CD45 endodomain fused to the Thy-1 ectodomain (Thy1-CD45*). All constructs contain an N-terminus Flag-Tag (shown in red). **b.** Fluorescence histograms of cell-surface levels of the indicated CD45 chimeras in sorted G8 T cell hybridomas transductants. CD43-CD45 transductants were labeled with secondary antibody only as a labeling control. **c. Top panel**, Timeseries of calcium flux fluorescence imaging of Fluo-4-loaded G8 T cell hybridomas, transduced with the indicated CD45 chimeras, interacting with SLBs containing 100 molecules/μm^2^ T22 and 350 molecules/μm^2^ ICAM-1. Images at each time point are centered averages of 25 cells for CD43-CD45, 20 cells for Thy1-CD45, and 30 cells for Thy1-CD45*. Pseudocolor scale indicates Fluo-4 fluorescence intensity in arbitrary units. **Bottom panel**, Quantitation of Flou-4 fluorescence intensity over time as a measure of intracellular Ca^2+^ flux in G8 T cell transductants. Data points are mean ± SEM; N=61 cells (CD43-CD45), 29 cells (Thy1-CD45), 51 cells (Thy1-CD45*), pooled from 4 independent experiments. **d.** IL-2 release by indicated G8 T cell transductants in 1:1 co-cultures with CHO cells expressing T22. Results are representative of 5 independent experiments. **e.** Schematic of wild-type CD148 (WT-CD148) and an ectodomain truncation retaining two membrane-proximal fibronectin type III domains (2FN3-CD148), depicted approximately to scale. Also shown is a chimera of a catalytically inactive version of 2FN3-CD148 (2FN3-CD148*). **f.** Fluorescence histograms of cell-surface expression levels of CD148 chimeras in stably transduced and sorted G8 T cell hybridomas expressing the indicated chimeras. **g. Top panel**, Timeseries of averaged epifluorescence images of Flou-4-labeled G8 T cell transductants expressing the indicated CD148 chimeras, interacting with SLBs as in ***c.***. Images at each time point are centered averages of 22 WT-CD148-, 18 2FN-CD148-, and 25 2FN-CD148dead-expressing G8 T cell hybridomas. **Bottom panel**, Quantitation of Flou-4 fluorescence intensity over time as in ***c.***. Data points are mean ± SEM. N=46 cells (WT-CD148), 40 cells (2FN-CD148), 38 cells (2FN-CD148dead), pooled from 2 independent experiments. **h.** IL-2 release by indicated G8 T cell transductants in 1:1 co-cultures with CHO cells expressing T22. Results are representative of 5 independent experiments. ****, P < 0.0001; ***, P < 0.001; *, P < 0.05; ns, not significant (P > 0.05); P values are corrected for all pairwise comparisons. Statistical comparisons in ***c*** and ***g*** are for data points at T=120s.

We measured ligand-induced Ca^2+^-flux in transduced G8 T to determine if truncating the CD45 ectodomain affected TCR triggering. T cell transductants were loaded with Fluo-4 fluorescent Ca^2+^ indicator, and intracellular Ca^2+^ influx monitored by live-cell epifluorescence microscopy of T cells interacting with SLBs containing T22. G8 T cells transduced with CD43-CD45 exhibited robust Ca^2+^ influx within 30 seconds of contact with bilayers (Fig. 5c). Strikingly, Ca^2+^ influx was abolished in G8 T cell hybridomas expressing Thy1-CD45 at 0.5% of endogenous CD45 levels, but was unaffected by expression of comparable levels of catalytically inactive Thy1-CD45 (Fig. 5c, Supplementary Table 1), indicating that CD45-mediated dephosphorylation was responsible for the abrogation of triggering by Thy-1-CD45. Consistent with this, IL-2 release by Thy1-CD45-expressing T cell transductants was significantly reduced in cocultures with T22-expressing CHO cells, but was unaffected in Thy1-CD45*-expressing G8 T cells (Fig. 5d).

Using a similar approach, we next generated stable transductants of G8 T cell hybridomas expressing comparable surface levels of full-length CD148, a truncated form containing 2 (of 7) of its fibronectin 3 domains (2FN3-CD148), and a truncated construct that was catalytically inactive (2FN3-CD148*) (Fig. 5e,f)^35^. Overexpression of wild-type CD148 in G8 T cell hybridomas stimulated robust Ca^2+^responses to T22 on SLBs, while Ca^2+^-influx was abrogated in transductants expressing 2FN3-CD148, which was partially restored by inactivation of its phosphatase activity (2FN3-CD148*) (Fig. 5g). In keeping with the observed Ca^2+^ responses, 2FN3-CD148 expression substantially reduced IL-2 release, which was fully restored in G8 T cell hybridomas expressing 2FN3-CD148* (Fig. 5h). Taken together, these results demonstrate that the large ectodomains of CD45 and CD148 are necessary for ligand-induced γδ TCR phosphorylation, and suggests that truncating their ectodomains abrogates segregation of from TCR at synaptic contacts.

### CD45 ectodomain size controls nanoscale segregation from T22-engaged G8 TCRs

To investigate whether ectodomain size controls CD45 segregation from TCR at synaptic contacts, we performed two-dimensional *direct* stochastic optical reconstruction microscopy (*d*STORM) using a TIRF (total internal reflection fluorescence) illumination platform^36, 37^. TIRF excitation substantially improved imaging signal/noise and localization accuracy, resulting in global image resolution of ∼55 nm as estimated by Fourier ring correlation^38^. TCR and CD45/Flag-tag constructs were labelled as for confocal microscopy, using secondary F(ab’)_2_ fragments conjugated with defined numbers of Atto488 and AF647 fluorophores, which spontaneously photoswitch under mild reducing conditions^39^. G8 T cell hybridomas/transductants were fixed after 2 minutes incubation at 37°C with T22-containing SLBs, and processed for two-channel *d*STORM imaging (Fig. 6a,b). Localizations were mapped and analyzed using Ripley’s H function L(r)-r^40, 41^ for single-channel 2D spatial point autocorrelation (amplitude indicates relative cluster density and *r* at maxima indicates approximate cluster radius) and two-channel crosscorrelation (negative values indicate segregation). TCRs were highly clustered at G8 T cell synapses, with a predominant cluster diameter of ∼150nm (Fig. 6c). In contrast, endogenous CD45 exhibited minimal clustering, with an autocorrelation amplitude and cluster size comparable to fluorophore localizations of primary and secondary antibody complexes alone (Fig. 6c, Extended Data Fig. 5). Crosscorrelation analysis revealed CD45 segregation from TCR at nanometer length scales (>200 nm), consistent with CD45 segregation from TCR clusters (Fig. 6d). Comparably-sized CD43-CD45 also exhibited nanoscale segregation from TCR (Fig. 6e,f). In stark contrast, CD45 chimeras with Thy-1-substituted ectodomains, Thy1-CD45 and catalytically inactive Thy1-CD45*, were no longer segregated from TCR, and could be detected within TCR clusters (Fig. 6e,f). These results demonstrate that ectodomain size is the principal determinant of CD45 segregation from TCRs at synaptic contacts. The large ectodomain of endogenous CD45 is required for segregation from TCR, which is preserved when the CD45 ectodomain is substituted with the comparably sized but structurally unrelated ectodomain of CD43. Segregation from TCR is abolished by substitution of the CD45 ectodomain with the small single IgSF Thy-1 ectodomain.

**Figure 6.**
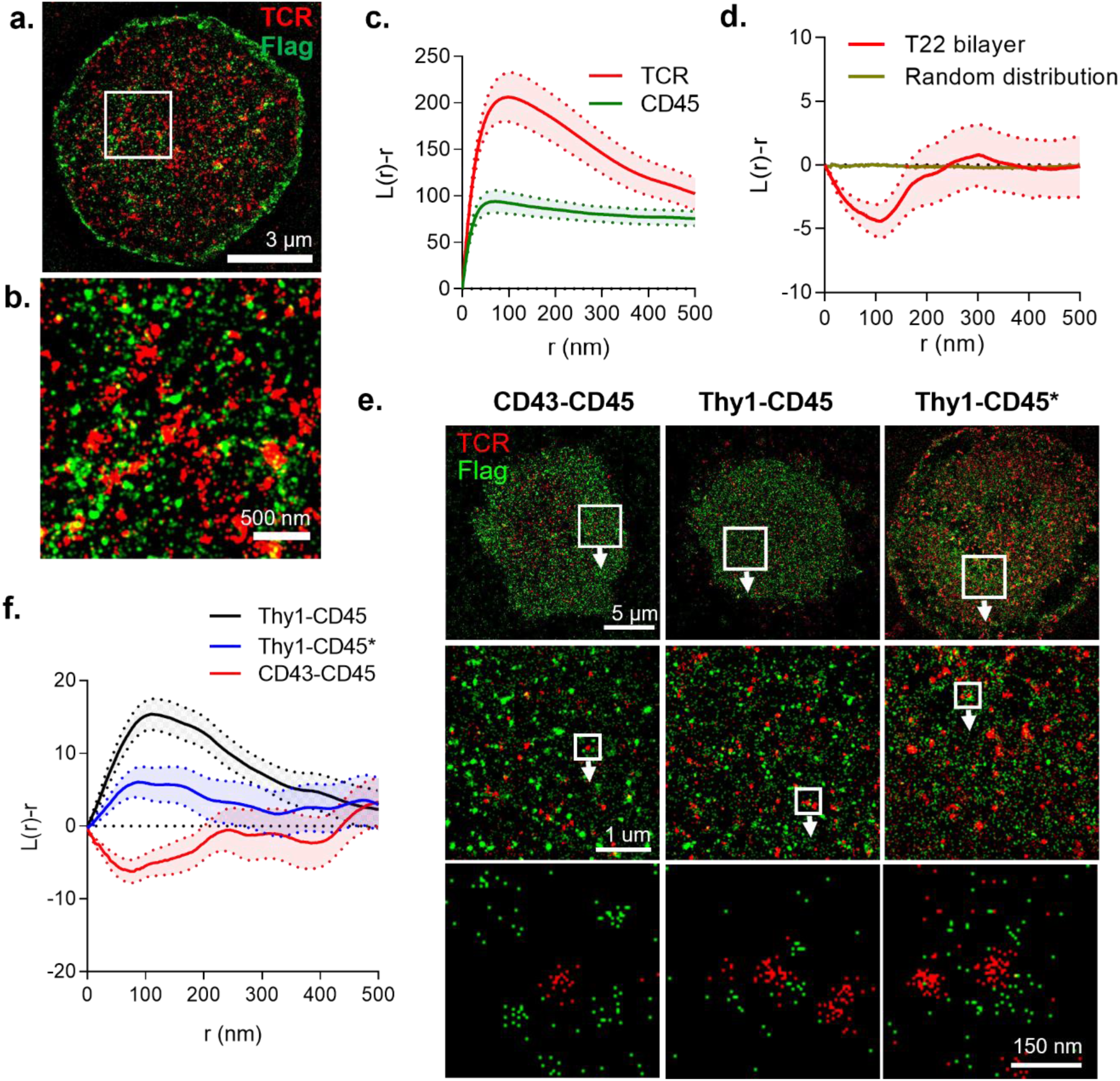
Truncated CD45 fails to segregate from γδ TCR at synaptic contacts. **a**. Representative image of a TIRF-*d*STORM Gaussian-filtered overlay of endogenous TCR and CD45 localizations at a G8 T cell synapse, formed on a SLB containing 100 molecules/μm^2^ T22 and 350 molecules/μm^2^ ICAM-1. Image intensity approximates localization density. **b.** High magnification image of boxed region in ***a.***. **c.** Autocorrelation of TCR and CD45 localizations as a function of distance calculated as Ripley’s H function. Peak values approximate cluster size (nm) and extent of clustering. Graphs represent mean ± SEM of 14 ROIs from 14 synapses of G8 T cells pooled from 3 independent experiments. Shaded envelope within dotted lines represents ± SEM. **d.** Cross-correlation of endogenous TCR and CD45 using Ripley’s H function. Negative values indicating dispersion (segregation), and positive values indicate co-clustering. Random distribution represents the mean 2D Poisson distributions from 20 Monte-Carlo simulations. **e.** G8 T cell hybridoma transductants forming synapses with SLB as in ***a.***; **Top panels,** Representative TIRFM-STORM Gaussian-filtered overlays of TCR and the indicated CD45 chimeras (labeled using an anti-Flag or anti-CD3ε antibody and appropriate fluorescently-labeled F(ab’)_2_ secondary; **middle panels**, High magnification images of boxed region in top panels; **Bottom panels**, Unfiltered localizations of TCR and CD45 chimeras in boxed regions of middle panels (each dot represents a single fluorophore localization). **f.** Cross-correlation of TCR and CD45 chimeras in G8 T cell transductant forming synapses with T22-containing supported bilayers. Graphs show mean ± SEM. N=29 ROIs from 13 cells (CD43-CD148), 38 ROIs from 13 cells (Thy1-CD148), 84 ROIs from 13 cells (Thy1-CD45*), pooled from 3 independent experiments.

### Triggering of sulfatide/CD1d-specific human γδ TCR DP10.7 requires CD45 segregation

We next sought to determine whether phosphatase segregation was also a critical requirement for human γδ TCR triggering. As in mice, nonclassical MHC class Ib molecules are a prominent class of antigens recognized by human γδ T cells, primarily of the Vδ1+ subset^42, 43, 44^. Since most of the identified MHC class Ib ligands recognized by Vδ1+ T cells are lipid-presenting molecules, we chose to focus on the well-characterized Vγ4Vδ1 TCR DP10.7, which recognizes sulfatide lipids presented by the nonclassical MHC I molecule CD1d^45^. We expressed a previously described^45^ CD1d SCD construct in CHO cells, and used this as a template to generate elongated versions of CD1d by insertion of the 2 IgSF ectodomain of mouse CD2 (CD1d-CD2) and the 4 IgSF ectodomain of mouse CD4 (CD1d-CD4) into the CD1d SCD membrane-proximal stalk (Fig. 7a). CHO cell transductants were sorted for comparable cell surface expression levels and exogenously loaded with soluble sulfatide lipid (24:1 acyl chain). We confirmed comparable sulfatide loading of CD1d constructs by labelling cells with a fluorescently-tagged DP10.7 TCR tetramer, and analysed surface staining by flow-cytometry (Fig. 7b). DP10.7 TCR triggering in response to sulfatide-loaded CD1d constructs was assessed using a TCR β-chain-deficient human Jurkat T cells (J.RT3-T3.5) stably transduced with DP10.7 TCR (DP10.7 Jurkat T cells), as described previously^45^. As with murine γδ TCRs, the binding-induced conformational change in CD3ε was absent following engagement of DP10.7 TCR by cell-expressed sulfatide/CD1d or by soluble sulfatide-loaded CD1d tetramers, as assessed by Nck SH3.1 pulldown and TCRζ immunoblot (Fig. 7d). Despite this, DP10.7 TCR was robustly phosphorylated in response to cell-expressed sulfatide-CD1d, resulting in downstream LAT phosphorylation (Fig. 7e), and T cells activation as measured by CD69 upregulation (Fig. 7f). However, TCR triggering and T cell activation was drastically attenuated by CD1d elongation, demonstrating that the small size of CD1d was critically important for effective TCR triggering (Fig. 7e,f). This was not due to impaired TCR engagement, as DP10.7 Jurkat T cells formed comparable levels of antigen-induced conjugates with CHO cells presenting sulfatide-loaded CD1d and elongated forms of CD1d, as measured by a flow-cytometry-based adhesion assay (Fig. 7c).

**Figure 7.**
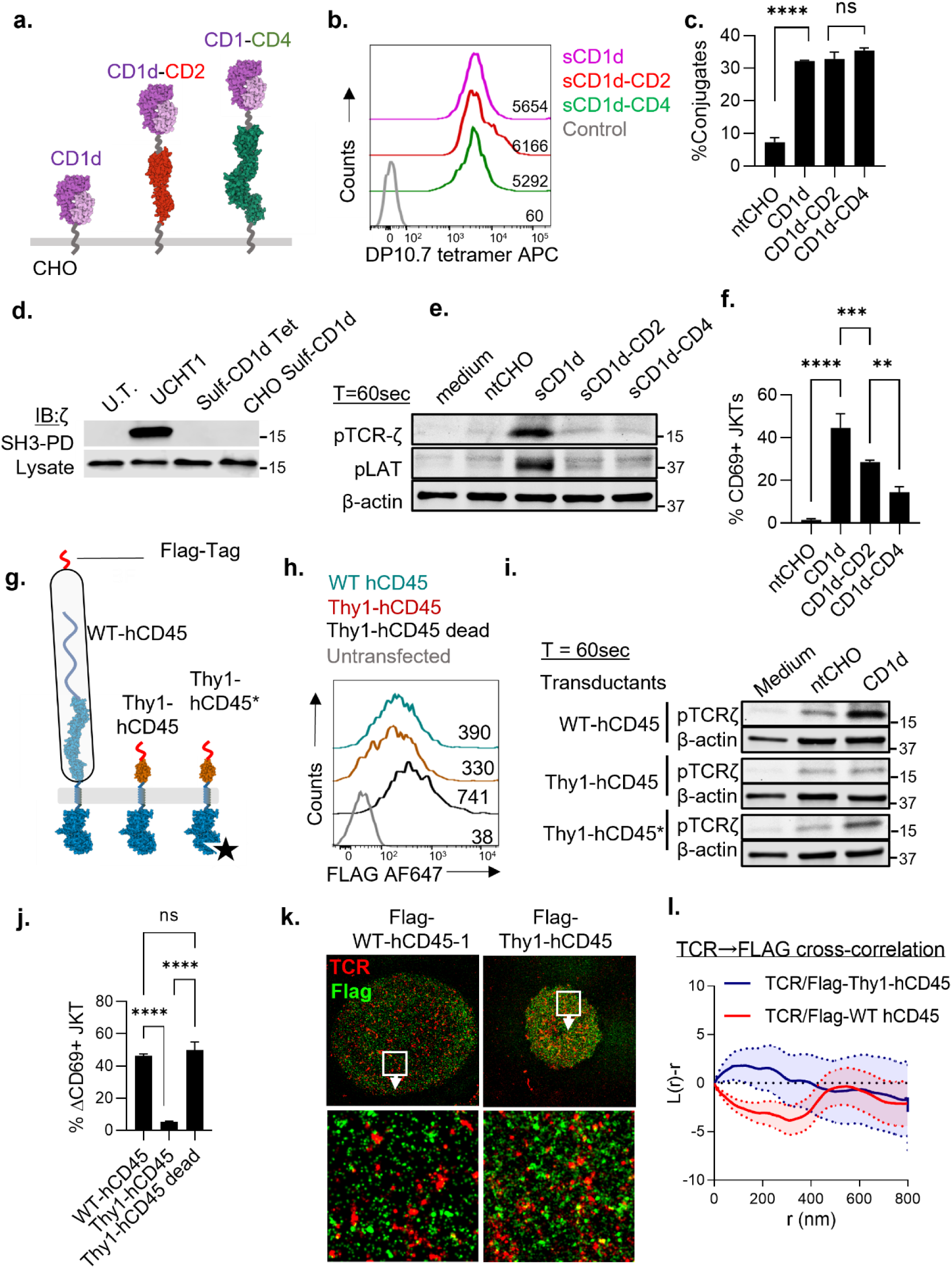
Triggering of human γδ TCR DP10.7 by sulfatide-CD1d requires phosphatase segregation. **a**. Schematic of single-chain constructs of human CD1d and elongated forms of CD1d incorporating membrane-proximal mouse CD2 and CD4 spacers. **b.** Fluorescence histograms showing cell-surface expression levels of sulfatide-loaded CD1d chimeras in CHO cell transductants. Sulfatide/CD1d complexes were detected with cognate DP10.7 TCR tetramer conjugated with APC. Results are representative of 4 independent experiments. **c.** Conjugate formation, as a measure of cell-cell adhesion, between fluorescently labeled Jurkat T cells expressing the sulfatide/CD1d-specific TCR DP10.7 (CMFDA) and CHO cells expressing the indicated CD1d chimeras loaded with sulfatide (Celltracker Deep Red). Data are pooled from 3 independent experiments. Data shown are mean+SD. **d.** Detection of CD3ε conformational change by Nck SH3.3 TCR pulldown in DP10.7 Jurkat T cells. Cells were incubated with anti-CD3ε mAb UCHT1, sulfatide-loaded CD1d tetramer, or left untreated (U.T.). **e.** Immunoblot of indicated tyrosine-phosphorylated TCR proximal signaling components (pTCR-ζ and pLAT) following 60 seconds contact at 37°C with CHO cells expressing the indicated CD1d constructs. Results are representative of 6 independent experiments. **f.** Activation of DP10.7 Jurkat T cells co-cultured with CHO cells expressing the indicated constructs, depicted as percentage change in CD69 levels in relative to baseline, as measured by flow-cytometry. Results are representative of 5 independent experiments. **g.** Schematic of expression constructs of wild-type human CD45 (WT-hCD45) and a chimera of the CD45 endodomain fused to the ectodomain of rat Thy-1 (Thy1-hCD45), depicted approximately to scale. Also shown is a chimera of a catalytically inactive human CD45 endodomain mutant fused to the Thy-1 ectodomain (Thy1-hCD45*). All constructs contain an N-terminal Flag-Tag. **h.** Fluorescence histograms of cell-surface expression levels of the indicated CD45 chimeras in DP10.7 Jurkat T cell transductants sorted for comparable surface expression. **i.** Immunoblot of tyrosine-phosphorylated TCRζ and LAT in DP10.7 Jurkat T cells after 60 seconds contact with CHO cells presenting the indicated sulfatide-loaded CD1d constructs. Representative of 2 independent experiments. **j.** CD69 upregulation, as measured by flow-cytometry, of DP10.7 Jurkat T cells expressing the indicated hCD45 constructs, in co-culture with CHO cells presenting the indicated sulfatide-loaded CD1d constructs. Results are representative of 2 independent experiments. **k. Top panels**, representative Guassian-filtered *d*STORM overlay images of endogenous TCR and indicated CD45 chimeras at synapses formed by the indicated Jurkat T cell transductants on supported lipid bilayers containing 100 molecules/μm^2^ sulfatide-loaded CD1d and 350 molecules/μm^2^ ICAM-1; **Bottom panels**, High magnification images of boxed region in ***top panels***; **l.** Ripley’s H cross-correlation between endogenous TCR and transduced CD45 chimeras at synapses of indicated DP10.7 Jurkat T cell transductants. Data are mean±SEM of 93 ROIs from 15 WT-CD45 transductants, and 33 ROIs from 11 Thy1-CD45 transductants, pooled from 3 independent experiments. ****, P < 0.0001; ***, P < 0.001; **, P < 0.01; ns, not significant (P > 0.05); P values are corrected for all pairwise comparisons.

To assess whether the size of the CD45 ectodomain affected human γδ TCR triggering, we made a human CD45 chimera, similar in design to our mouse CD45 constructs, in which the native CD45 ectodomain was replaced with that of rat Thy1 (Thy1-hCD45) (Fig. 7g). We also generated expression constructs for full-length native human CD45 (WT-hCD45) and a phosphatase-inactivated mutant of Thy1-hCD45 (Thy1-hCD45*) (Fig. 7g). All constructs contained an N-terminal Flag tag for sorting and immunodetection (Fig. 7h). DP10.7 Jurkat T cell transductants that stably expressed CD45 constructs were labelled with an anti-Flag antibody and sorted for comparable surface expression levels. TCR triggering and DP10.7 Jurkat T cell activation, as measured by TCRζ phosphorylation (Fig. 7i) and CD69 upregulation (Fig. 7j) respectively, was unaffected by expression of hCD45 at ∼6% of endogenous CD45 levels (Supplementary Table 1). In marked contrast, TCR triggering and T cell activation was almost completely abrogated in DP10.7 Jurkat T cells expressing the truncated Thy1-hCD45 construct at ∼5% of endogenous CD45 levels (Fig. 7i,j and Supplementary Table 1). In contrast, TCR triggering and T cell activation was unaffected by catalytically inactive Thy1-hCD45*, despite expression at more than twice the levels of Thy1-hCD45 (∼11% of endogenous CD45). Like endogenous CD45, transduced WT-hCD45 remained segregated from TCR at DP10.7 Jurkat T cell synapes with CD1d-sulfatide-containing SLBs, as determined by *d*STORM imaging (Fig. 7k). Spatial point cross-correlation analysis of TCR/hCD45 localizations using Ripley’s H function revealed negative values at distances < ∼400 nm, indicating lateral segregation of WT-hCD45 from TCR at receptor-level length scales (Fig. 7l). In contrast, Ripley’s H function for TCR/Thy1-hCD45 approximated zero at all length scales, representing a loss of TCR/CD45 segregation (Fig. 7l). Taken together, these results demonstrate that the large ectodomain of human CD45 is necessary for effective segregation from engaged DP10.7 TCR, and is required for stable ligand-induced TCR phosphorylation at synaptic contacts.

## Discussion

Here, we identify size-based steric segregation of engaged TCRs from large T cell surface tyrosine phosphatases CD45 and CD148 as a critical requirement for γδ TCR triggering. This is in keeping with the K-S model of TCR triggering^15^, which does not depend on binding-induced mechanical forces, conformational changes, or coreceptor-mediated aggregation, but only requires that membrane-tethered cognate ligands are small in size relative to the large ectodomains of T cell surface tyrosine phosphatases.

We show, in two well characterized γδ TCR/ligand pairs, murine G8/T22^17^ and human DP10.7/sulfatide-CD1d^45^, that increasing phosphatase access to engaged TCRs abrogates triggering. Engagement of the G8 γδ TCR by elongated T22 increases intermembrane distance of apposed plasma membranes at T cell synaptic contacts. Despite effective TCR engagement by elongated T22, loss of close-contacts abolishes CD45 segregation from engaged TCRs, and abrogates TCR triggering. While we show that ligand elongation does not affect overall TCR binding, it remained possible that engagement by elongated ligands held TCRs in orientations unfavourable for triggering. To address this, we increased the flexibility of elongated T22 to enable greater degrees of freedom for TCR engagement. This did not rescue TCR triggering, as would be expected if TCR binding were aberrantly constrained in unfavourable orientations. Moreover, we show that γδ TCR triggering in response to native-sized T22 and sulfatide-CD1d is abrogated by expression of ectodomain-truncated phosphatases in T cells, where the TCR-ligand interaction is unperturbed, hence ruling out inadvertent disruption of binding-induced mechanical forces or conformational changes. Our findings, however, do not preclude the involvement of conformational changes or mechanical forces in γδ TCR signal transduction. We envisage that such processes would complement the core K-S triggering mechanism that we identify here, for instance, by increasing antigen sensitivity through amplifying TCR-evoked proximal signals, or by enhancing antigen specificity by sensing binding-induced force signatures of cognate ligands^46^.

Interestingly, in contrast to αβ T cells, artificial aggregation of G8 or DP10.7 TCRs by soluble T22 or sulfatide-CD1d tetramers fails to activate γδ T cells, suggesting that ligand-induced oligomerization alone, in the absence of close contacts, is insufficient to trigger γδ TCRs. Strict reliance on the K-S mechanism for γδ T cell antigen sensing, especially in the absence of coreceptor-mediated signal amplification^47^, likely reflects a higher signaling threshold for γδ T cell activation. This may be important for calibrating γδ T cell antigen sensitivity to the relatively higher levels of basal MHC class Ib ligands on cell surfaces in comparison to the handful of cognate pMHC ligands typically displayed on antigen presenting cells. Similarly, our finding that Csk downmodulation had no impact on γδ TCR triggering suggests that stress-induced upregulation of abundant nonpolymorphic ligands on cell surfaces obviates the need for Csk-mediated fine-tuning of γδ TCR signaling thresholds to adjust for variability in ligand potency^29^.

Structural studies have revealed substantial variability in γδ TCR docking angles and ligand-binding topology, in contrast to the stereotypical end-to-end orientation and ‘diagonal’ binding mode of αβ TCRs to cognate pMHC ligands^48^. Both the murine G8 TCR and the human DP10.7 TCR bind their MHC class Ib ligands predominantly with their δ-chains^17, 45^, while some MR1-restricted γδ TCRs bind to the MR1 α3 domain, below the antigen presenting platform^49^. This surprising flexibility in ligand binding, free from the constraints imposed by peptide recognition or coreceptor engagement, suggests that effective γδ TCR triggering is indifferent to the site and orientation of binding to cognate ligands, as long as TCR engagement takes place in synaptic close contact zones^50^, from which phosphatases are excluded. This B cell receptor-like recognition of membrane-tethered invariant antigens is reminiscent of antigen targeting by chimeric antigen receptors (CAR). CARs employ small antibody fragments for targeting extracellular tumor antigens, fused to ITAM-containing cytoplasmic domains^51^, fulfilling conditions for triggering according to K-S principles. In support of this, CAR triggering has been recently been shown to be dependent on compact CAR and antigen dimensions^52^.

Finally, our findings raise the possibility that the K-S mechanism may also be involved in TCR triggering in human Vγ9Vδ2 T cells. These γδ T cells are activated by exposure to phosphorylated isoprenoid metabolites (PiPs)^4^, although the mechanism by which PiPs activate Vγ9Vδ2 T cells remains unclear. Vγ9Vδ2+ T cells require contact with accessory cells for activation^53^, and CD45 is depleted at synaptic contacts^54^. Recently, the small cell surface glycoprotein BTN2A1, a member of the B7 receptor family, was identified as a ligand for Vγ9Vδ2+ T cells^55, 56^. BTN2A1, which possesses a small 2 IgSF ectodomain, directly engages the TCR and may therefore mediate TCR triggering, at least in part, by trapping TCR in close contact zones from which T cell surface phosphatases have been excluded.

## Acknowledgements

We thank YH Chien (Stanford University) for the generous gift of mAb 7H9, G8 hybridoma and the *tcrG8rag-/-Balb/c* mice, and Luc Van Kaer (Vanderbilt University) for kindly providing KN6 hybridomas. This work was supported by NIH grants K99AI093884, R00AI093884 and R01AI134999 to K.C., grant NIH R01AI155984 to E.J.A, and an MRC Programme Grant (G9722488) and Wellcome Trust Senior Investigator Award (101799/Z/13/Z) to P.A.v.d.M. We thank the University of Michigan Biomedical Research Core Facilities and Rogel Cancer Center (NCI award P30CA046592) for support with microscopy and flow-cytometry/cell-sorting resources. The following reagent(s) was/were obtained through the NIH Tetramer Core Facility: T22 and sulfatide/CD1d tetramers, and monobiotinylated T22 and biotinylated CD1d monomers.

## Author contributions

K.C and P.A.v.d.M. conceived of the study, K.C and P.A.v.d.M designed the study, F.L., K.C. and J.N. performed experiments, K.G. made T22 constructs, E.J.A. and S.R. made critical reagents and advised on study design for CD1d experiments, K.C. and F.L. wrote the manuscript draft, all authors contributed to editing the final manuscript.

## Data availability statement

All data associated with the present study are available from the corresponding authors upon request.

## Methods

### Recombinant expression constructs and siRNA vectors

T22 mRNA was isolated from LPS-stimulate C57BL/6 splenocytes. After RNA extraction, T22 cDNA was obtained using the Qiagen one-step RT-PCR kit according to the manufacturer’s instructions. T22, and β2-microglobulin (β2M) were amplified by PCR, using primers listed in Supplementary Table 2., such that β2M contained a 5’ XhoI site, T22 a 3’HindIII site while the primers for the β2M 3’ and T22 5’-ends together encoded for a (GGGGS)_3_ flexible linker with 20 base pair overlap between the two primers. Overlap extension PCR was then used to produce a single chain dimer (SCD) of T22, consisting of murine β2m (including the leader sequence), followed by a (GGGGS)_3_ linker and the complete T22 sequence. After digestion with HindIII and XhoI, this was then cloned into the pKG5 expression vector for transfection.

The elongated SCD chimeras were generated by first introducing a BamHI site into the extracellular stalk region by site-directed mutagenesis (using oligonucleotides listed in Supplementary Table 2), thereby changing the amino acid sequence by 1 residue, from the native WE**P**AW to WE**D**PAW. Human cDNA templates were used to create PCR products flanked with the BamHI restriction sites and encoding either the human CD2 ectodomain (amino acids 1-180), or human CD4 ectodomain (amino-acids 1-363). The elongated SCD constructs were then produced by inserting CD2 or CD4 into the BamHI site. For the T22 SCD constructs containing GGGGS or (GGGGS)_2_ flexible linkers, complementary 5’-phosphorylated oligonucleotides were annealed. The sequences of these oligonucleotides were such as to produce double-stranded DNA with 5’-overhangs of identical sequence to those produced by BamHI. These were then annealed into the T22 SCD BamHI site. Alternatively, 6 amino-acids of the wild type T22 stalk sequence after the BamHI site were first deleted by oligonucleotide-directed mutagenesis (using oligonucleotides listed in Supplementary Table 2) and converting WEDPAWYQDPWIW to WEDPWIW followed by insertion of the annealing oligos encoding the GGGGS or (GGGGS)_2_ flexible linkers. Finally, the T22-CD4 chimera was also mutated by site-directed mutagenesis (using oligonucleotides listed in Supplementary Table 2) to remove the membrane proximal BamHI site, allowing insertion of the annealing oligonucleotides encoding the GGGGS or (GGGGS)_2_ flexible linkers at the remaining membrane-distal BamHI site.

A single chain construct of human CD1d was generated, with a similar design to T22 SCD. The Gp64 signal peptide, full-length human β2M and a (G_4_S)_2_ linker were amplified from plasmid CD1C-NO403^57^ by PCR. hCD1d α chain ectodomain (aa24-295) was amplified from plasmid CD1d-590^58^ by PCR, while a segment comprising the hCD1d α chain transmembrane and cytoplasmic domains (aa296-335) were synthesized as dsDNA gBlock fragments (IDT). These three segments were ligated sequentially by overlap PCR and inserted into XhoI/NotI linearized Phr-SIN vector by In-fusion cloning (Clontech) to produce full length CD1d SCD. To produce elongated CD1d constructs, the CD1d transmembrane and cytoplasmic segment was first 5’-ligated with mouse CD2 (aa23-202) gBlock or CD4 (aa28-391) gBlock, then ligated with the other two upstream segments (GP64 signal peptide-human β2M-(G_4_S)_2_ linker and hCD1d α ectodomain) by overlap PCR, and the resulting CD1d-CD2 and CD1d-CD4 SCDs were cloned into Phr-SIN expression vector (as for CD1d SCD) for lentiviral transduction of CHO cells. Primers and gBlocks are listed in Supplementary Table 2.

Expression constructs for Thy1-CD45, Thy1-CD45* (*; inactivating mutation in the phosphatase domain), CD43-CD45, wtCD148, 2FN3-CD148 and 2FN3-CD148* are described in detail elsewhere^35^. The ectodomains are derived from rat Thy1, and human CD148 and CD43, and the phosphatase-containing intracellular domain of all constructs are derived from mouse CD45. Constructs were amplified from their original vectors by PCR, and cloned into Phr-SIN vector, using XhoI and NotI restriction sites, for lentiviral transduction. The C-terminus GFP sequence of the original constructs resulted in unstable expression in G8 T cell hybridomas, and was therefore removed by insertion of a TAG stop codon after the 3^rd^ amino acid residue in the 5’-end of the sequence encoding GFP (using mutagenesis oligonucleotides listed in Supplementary Table 2).

Human wild-type CD45 (WT-hCD45), Thy1-hCD45 and Thy1-hCD45* were made using a similar design to mouse CD45 chimeras. A human CD45 DNA template was amplified by PCR from Jurkat T cell cDNA using the primers listed in Supplementary Table 2. The CD45 signal peptide (aa1-25), Flag-tag, and full-length CD45 (aa26-1306), were amplified by PCR primers containing overlapping sequences, and ligated sequentially with XhoI/NotI-linearized Phr-SIN vector by In-fusion cloning. For the Thy1-CD45 expression construct, a rat Thy1 (aa20-135) gBlock and an overlapping CD45 segment containing the stalk/transmembrane/cytoplasmic domain of human CD45 (aa571-1306) were ligated by overlap PCR to replace full length CD45 ectodomain. The Thy1-CD45* construct was generated by site-directed mutagenesis by C853S substitution. Primers and gBlocks are listed in Supplementary Table 2.

For Csk knockdown, SMARTvector lentiviral shRNA expression vectors encoding targeting and control (non-targeting) sequences listed in Supplementary Table 2 under the CMV promoter were purchased from Dharmacon and used according to the manufacturer’s instructions.

### Cell transfection and transduction

TAP2 deficient CHO cells were cultured in complete RPMI (Invitrogen, 2mM glutamine and 10% FCS) and stably-transfected with >10g plasmid DNA using poly-L-ornithine and a 30% DMSO shock, as previously described^24^. Stable transfectants were selected using 0.8mg/ml geneticin (Invitrogen) and maintained as polyclonal populations. For lentiviral transduction, HEK293T cells were transfected with constructs in Phr-SIN expression vectors, pRSV, pGAG, and pVSV-G using lipofectamine 3000 (ThermoFisher), to make lentiviral particles for transduction of G8, Jurkat and CHO cells. Matched expression levels was checked by flow-cytometry using Armenian-hamster 7H9 anti-T22 (gift from YH Chien, Stanford) for T22-expressing CHO, anti-Flag-tag M2 for Flag-CD45/CD148-expressing G8 and Jurkat T cells, and anti-CD1d antibody for CD1d-expressing CHO cells.

For Csk knockdown, HEK293T cells were transfected with shRNA lentiviral expression vectors containing *CSK*-targeting shRNA or a control non-targeting RNA sequence derived from firefly luciferase (Supplementary Table 2), along with pMD2.G and psPAX2 vectors. Lentiviral particles were harvested and used to transduce G8 T cell hybridomas. Transduced G8 cells were selected using puromycin and monitored by RFP expression. Lentiviral transduction was typically performed by ‘spinduction’ at 1000g for 1hr at 32°C in neat RPMI containing 5µg/ml polybrene. Following 24hrs incubation at 37°C, 5% CO_2_, cells were washed and maintained in complete RPMI.

### Flow-cytometry and cell sorting

To label cells for flow-cytometry and cell sorting, live cells (5 x 10^5^ cells/ml) were suspended in PBS/2% FBS buffer, Fc-blocked as appropriate, and stained with surface florescence antibodies on ice. The stained cells were washed and filtered through a 70µM cell strainer before analysis using an LSRFortessa (BD) flow-cytometer. Cell sorting was performed using FACSAria-Ι (BD) or Sony SH800 instruments fitted with a 100µm nozzle (FacsAria-Ι) or 100µm chip (SH800). Fluorescently labelled cells were gated to select for singlets using FSC-H/FSC-W and SSC-H/SSC-W channels, and a serial fluorescence gate used to sort cells with defined expression levels of labelled cell surface proteins. Cells were sorted into tubes containing cold FBS, and transferred to complete RPMI to rest and expand for 3-5 days prior to use in experiments. Sorted cells were checked for matched levels post-sort, and prior to experiments. Flow cytometry data analysis was performed using Flowjo10 software.

### Mice

A breeding pair of mice that transgenically express the G8 γδ TCR (bred on a Balb/c Rag^-/-^ genetic background) were a gift from YH Chien (Stanford). Colonies were maintained under specific pathogen-free (SPF) housing conditions at the University of Michigan Medical School animal housing facility, under an Institutional Animal Care & Use Committee (IACUC) approved protocol (PRO00008433). Mice were maintained in accordance with local, state and federal regulations. 8–10-week-old male and female mice were used for experiments. Both male and female mice were used in experiments.

### Cell isolation and enrichment

Primary G8 T cells were isolated by negative selection, from spleens harvested from G8 TCR transgenic mice, using an EasySep™ Mouse T Cell Isolation Kit (Stemcell Technologies) according to the manufacturer’s instructions. Briefly, splenocytes were prepared as a single-cell suspension by maceration and straining through a 45μm cell strainer. Rat serum and cell separation antibody cocktails and associated magnetic beads were added, and labelled cells were removed by magnetic isolation (Big Easy, Stemcell). Enriched G8 T cells were checked for purity by flow cytometry and rested overnight in complete RPMI media before experiment. Cell purity was checked by labelling with fluorescently-labeled anti-mouse CD3ε antibody and T22 tetramer (NIH Tetramer Core Facility).

### DP10.7 tetramer production

DP10.7 TCR was expressed in insect cells, biotinylated and tetramerized as reported previously^45^. Both chains were co-expressed in Hi5 cells by baculovirus transduction. The heterodimeric TCR was purified from cell culture supernatant with Ni-NTA Agarose (Qiagen) and treated with 3C protease to remove tags. TCR was biotinylated with BirA biotin protein ligase in the presence of biotin, and purified over an S200 size exclusion column. TCR was tetramerized with streptavidin-allophycocyanin (Agilent).

### T cell activation assay

For murine T cells and hybridomas, activation assays were performed as previously described^24^. Briefly, for G8 T cell hybridomas, cells (10^4^/well) were incubated with varying numbers of T22-expressing CHO cells (shown as APC/G8 ratio) at 37°C in 5% CO_2_ for 24 hours. In experiments using G8 T cell hybridoma transductants expressing CD45 and CD148 chimeras, 10^4^ cells/well were co-cultured with equal numbers of CHO cells. Culture supernatants were assayed for IL-2 by sandwich ELISA using 1µg/ml capture antibody (JES6-1A12), and 1µg/ml biotinylated detecting antibody (JES6-5H4) followed by labelling with 1:2000 diluted ExtrAvidin®−Peroxidase (Sigma E2886) and developed with 1-Step™ Ultra TMB-ELISA Substrate Solution (Thermo Scientific™). For primary G8 T cells, experiments were performed as for T cell hybridomas, but mouse IFN-γ was assayed in culture supernatants using 4µg/ml capture antibody (R4-6A2), and 1µg/ml biotinylated detecting antibody (XMG1.2). For DP10.7 Jurkat T cells, CHO cells expressing CD1d constructs (10^5^/ml) were incubated with 30ug/ml sulfatide (Matreya LLC) in complete RPMI for 1hr at 37 degree 5% CO2, before washing and coculture with equal numbers of DP10. JKT cells. CD1d loading was measured using a CD1d-APC tetramer and flow-cytometry. Activation of DP10.7 Jurkat T cells and CD45 transductants was assayed by measuring CD69 upregulation (expressed as % change in mean surface CD69 levels) by flow cytometry (Extended Data Fig. 6b).

### Adhesion Assay

Experiments were performed as previously described^24^. Briefly, to assay T22-CHO/G8 adhesion, CHO and G8 hybridomas cells were loaded respectively with 10 μM of the red cell-labeling agent DDAO (Molecular Probes) [ex/em 635/662] and 1 μM Vybrant CFDA SE [ex/em 492/517] in serum-free RPMI-1640 for 30 minutes at 37°C. After 3 washes and 30 minutes 37°C incubation in complete media, equal numbers of CHO and G8 (2 x 10^5^) were mixed and centrifuged at 400rpm/4°C (brake off). Cells were incubated for 10min at 37°C gently resuspended, and immediately analysed using a flow-cytometer (FacScan, BD). To assay CD1d-CHO/JKT adhesion, CHO and JKT cells were loaded respectively with 2 μM CellTracker™ Deep Red [ex/em 630⁄650] and 0.5 μM CellTracker™ Green CMFDA [ex/em 492/517] for 30 minutes at 37°C, after 3 washes and 30 minutes 37°C incubation in complete media, CD1d-CHO were incubated with 30 μg/ml Sulfatide for 60 minutes at 37°C. equal numbers of CHO and JKT (2 x 10^5^) were mixed and centrifuged at 1000rpm/4°C (brake off). Cells were incubated for 10 minutes at 37°C, gently resuspended, and immediately analysed using a flow-cytometer (LSRFortessa, BD). Adhesion assays between CHO cells expressing comparable complexes of sulfatide-CD1d constructs (detected with DP10.7 tetramer) and DP10.7 Jurkat T cells were performed using the same protocol (Extended Data Fig. 6a).

### Biochemistry

Immunoprecipitation and immunoblotting for TCRζ was performed as previously described^24^. Briefly, equal numbers of G8 cells and CHO APCs expressing the indicated SCDs were rapidly brought into contact by pulse centrifugation at 500g for 1 minute at 4°C. After incubation at 37 °C for 0, 2, 5 or 10 min, cells were lysed with chilled lysis buffer (50 mM Hepes, pH7.4, 150 mM NaCl, 1% NP-40, 1 mM PMSF, 10 mM pervanadate, protease inhibitor cocktail (Sigma). For samples at the 0 time point, cells were lysed on ice immediately following centrifugation. To detect TCR conformations that bind Nck, a pulldown assay was designed based on a well-established protocol. Briefly, C-terminal mono-biotinylated Nck SH3.1 peptides were immobilised on streptavidin-conjugated magnetic beads (Pierce) and washed thoroughly. 10^7^ T cells were incubated at 37°C for 10 minutes with anti-CD3ε mAb (2C11), T22 tetramers, CHO cells expressing T22, or sulfatide loaded CD1d. Cells were lysed using RIPA buffer (0.3% Brij O10, 140mM NaCl, 50mM Hepes, 10mM iodoacetamide, 1% protease inhibitors, 1mM sodium metavanadate), and 10^7^ T cells derived lysate were incubated with 10^7^ Nck peptide-immobilized beads at 4°C on a rotating mixer for 4 hours. Beads were washed 5 times with RIPA buffer and eluted by boiling in loading buffer containing 20mM DTT. Samples were separated by SDS-PAGE, transferred to PVDF membranes, and immunoblotted for TCRζ using anti-CD3ζ mAb (51-6527GR), and IRDye® 680LT Donkey anti-Mouse IgG (LI-COR). Blots were recorded using Odyssey® Classic Infrared Imaging System.

### TCR downregulation assay

G8 T cell hybridomas and CHO cells expressing T22 and elongated forms of T22 were gently centrifuged at 4°C and incubated in complete RPMI for 0-60 minutes at 37°C in 5% CO_2_. As a control, G8 hybridomas were co-cultured with untransfected CHO cells. At indicated time points, cells are resuspended and fixed with PFA of a final concentration of 2% for 15 minutes. Cells were then washed, quenched, labelled with Alexa-488-anti-CD3ε mAb (17A2), and analysed by flow cytometer.

### Supported lipid bilayers

Supported lipid bilayers were formed by liposome deposition on piranha-cleaned glass coverslips in Bioptechs flow chambers as previously described^59^. The resulting planar bilayers contained DOPC as the base lipid, and contained 12.5 mol% NTA-DGS (charged with Ni^3+^), and 0.1 mol% biotinyl Cap-PE. Recombinantly produced monobiotinylated T22 or CD1d alpha-chains complexed with β2M (obtained from the NIH Tetramer Core Facility) were attached to bilayers *via* a streptavidin bridge: SLBs formed in flow chambers were blocked with 4% casein containing 10mM NiSO_4_, and incubated with 5 mg/ml streptavidin (Thermo Scientific) for 15 minutes, followed by 15 minute incubation with monobiotinylated T22 or CD1d. Recombinantly produced ICAM-1-His10 (Sinobiologicals) was then attached to bilayers by coupling to Ni^3+^-charged NTA-DGS. Bilayers contained 100 molecules/μm^2^ of T22 or CD1d, and 350 molecules/μm^2^ of ICAM-1-10His. To load CD1d monomers with sulfatide, bilayers formed in flow chambers were incubated with 30ug/ml sulfatide in HBS for 1hr at 37°C, and sulfatide concentrations were maintained following introduction of T cells into flow chambers. Densities of T22, ICAM-1 and CD1d on SLBs were determined by interpolation from reference fluorophore beads as previously described^59^.

### Intracellular Ca^2+^ imaging and analysis

Live G8 T cell hybridomas, stably transduced with CD45/CD148 chimeras or shRNA expression vectors, were washed and resuspended in neat RPMI containing 2μM Fluo-4 (ex/em 494/506, Ca^2+^-bound) and incubated for 25 minutes at 37°C. Cells were then pelleted and resuspended in complete RPMI and rested for a further 30 minutes at 37°C. Equilibrated cells were resuspended at 3×10^6^/ml in HBS/HSA imaging buffer (20 mM Hepes, 137 mM NaCl, 5 mM KCL, 0.7 mM Na2HPO4, 6mM D-Glucose, 1mM CaCl2, 2mM MgCl2 and 1% HSA, PH 7.2), and injected into heated Bioptechs flow chambers containing SLBs containing T22 and ICAM-1-AF647. Fluo-4 fluorescence was imaged by wide-field microscopy (20x objective, 488nm laser excitation, and FITC emission filters) from the point cells settled on bilayers (T=0), and subsequently every 3 seconds for 2 minutes. 4 preselected imaging fields on each SLB were sampled sequentially at each timepoint using an encoded motorized stage with IR-based focus stabilization. The objective was pre-focussed on lipid bilayers using ICAM-AF647 fluorescence. Fluo-4 fluorescence was measured using 488nm laser excitation, 500-550nm emission filter (Nikon), and an EMCCD camera (Andor).

Quantitation of Fluo-4 fluorescence intensity was performed using Fiji/ImageJ by defining ROIs of cells contacting SLBs using a threshold above the image background. Mean fluorescence intensity of ROIs was plotted for all cells in an imaging field, starting with the image corresponding to T=0s (F_0_), the first frame in cell contact with bilayers was detectable, and in subsequent images of the same cell acquired every 3 seconds. As a measure of intracellular Ca^2+^ influx, the ratio of the difference in Fluo-4 fluorescence intensity from the initial F_0_ value (ΔF) over F_0_ (ΔF/ F_0_) was calculated for each cell over time, and mean values from all cells at each timepoint plotted as a time series.

### Confocal fluorescence microscopy

Conjugates of G8 primary T cells with CHO cells expressing T22 or T22-CD4 were made by mixing cells at 1:1 ratio in complete RPMI, and intercellular contact established by brief centrifugation at 4°C. Pelleted cells were then incubated at 37°C to initiate signaling. Following 2 minutes incubation, cells were fixed by replacing culture media with 3% PFA/PBS, gently resuspended, and left to settle on ice-cooled acid-washed coverslips coated with and poly-L-lysine for 20 minutes. Cells were permeabilized in 0.1% saponin/PBS for 5 minutes, quenched with 100 mM glycine/PBS, and blocked with 5% BSA/PBS. For labelling TCR and CD45 in G8 T Cells, samples were serially stained with 10μg/ml of anti-mCD3ε mAb 17A2 (1hr), followed by 10μg/ml of Atto-488 anti-rat F(ab’)_2_ fragments (45 min), 10ug/ml anti-mCD45.2 104 (1hr), and 10ug/ml AF647 anti-mouse F(ab’)_2_ fragments (45 min). Cells were washed thoroughly with 2% BSA/PBS after each labelling step. Labelled samples were mounted in ProLong™ Glass Antifade mountant (Invitrogen), and imaged using a NIKON A1+ inverted microscope fitted with a 60x oil objective (NA 1.4, Plan Apoλ, 60x). The pinhole was set to obtain ∼ 300 nm optical sections for each channel. AF647 was excited using a 640nm laser, and emission detected using 700/40nm emission filter, and Atto488 was excited with a 488nm laser and fluorescence images acquired using 525/50nm emission filter. DIC images were acquired using slider-mounted optics in the objective adapter. Images were recorded using and GaAsP PMT detector (Nikon). To acquire 3D image stacks, samples were imaged 200nm z-steps from the plane of the coverslip progressively into the sample. Synapse planes were isolated using the Reslice ImageJ plugin.

### *d*STORM sample preparation & imaging

For imaging G8 T cell hybridomas, 10^6^ T cells suspended in HBS/HSA were introduced into heated FCS2 flow chambers (Bioptechs) and allowed to interact with T22-containing SLBs as for *intracellular Ca^2+^-imaging*. After two minutes incubation, settled cells on SLBs were fixed with 3% PFA/HBS for 15 minutes. Flow-chambers were maintained at 37°C throughout using a filament heater attachment. Following fixation, cells were processed at room temperature as for *confocal fluorescence microscopy*. Endogenous murine TCR and CD45 were labelled with antibodies and secondary reagents as for confocal microscopy, and transduced Flag-tagged CD45/148 constructs were labelled by incubation with 10ug/ml anti-Flag-tag mAb M2 (1hr) followed by AF647-conjugated anti-mouse F(ab’)_2_ fragments. DP10.7 Jurkat T cell samples were prepared using a process. Cells were incubated in flow chambers with SLBs containing sulfatide-loaded CD1d and ICAM-1, then fixed and processed as for G8 T cell hybridomas. Endogenous TCR and CD45 in DP10.7 Jurkat T cells were labelled by serial staining with AF647-conjugated anti-hCD45 antibody H130 (1hr) and Atto-488-conjugated anti-hCD3ε antibody UCHT1 (1hr). Flag-tagged CD45 constructs in DP10.7 Jurkat T cell transductants were stained with 10ug/ml anti-Flag-tag polyclonal antibody (Sigma) for 1hr,

For *d*STORM imaging of T cell synapses on SLBs, HBS/HAS in flow chambers containing labelled samples was exchanged with STORM buffer (10 µM MEA, 0.56 mg/ml Glucose Oxidase, 40 µg/ml Catalase, 10 mM NaCl, 10% D-Glucose and 200 mM Tris-HCl (pH 8) and cells imaged on a Nikon TIRF-STORM inverted microscope fitted with a NIKON 100x TIRF objective (NA 1.49), using TIRF mode laser illumination. Laser outputs were 100% of a 70 mW 488 nm diode laser and 30% of a 125 mW 633nm diode laser with 0-5% of a 20 mW 405 mW diode laser, in line with a 4x beam magnification lens. Images were acquired with a Quad band N-STORM filter set (Nikon) fitted with excitation and emission filters for Atto-488 (ex/em: 488/505-545nm) and AF647 (ex/em: 647/664-787nm) and fluorescence detected using a sCMOS digital camera (Hamamatsu). Approximately 1,000 images were acquired alternately from each channel (total of ∼5000 images/channel), and assembled into two-channel image stacks for analysis.

### *d*STORM image processing

STORM image stacks were initially analysed using the ThunderSTORM^60^ localization plugin in Fiji/ImageJ^61^. Stacks were drift-corrected using the cross-correlation method, and background noise reduced by applying a B-spline wavelet image filter. Approximate positions of fluorophore emitters were localized by identifying local maxima of emitter spots, followed by Gaussian-fitting of spot PSFs for subpixel localization of the spot center using the maximum likelihood method and multi-emitter fitting. The resulting 2D localization data set was filtered to include fitted PSFs with a standard deviation of 50-250 nm, and a precision σ (√(SD^2^/photon count)) cut-off of ≤ 16 nm. This cut-off was empirically determined to be the minimum value that did not change the overall distribution of localization standard deviations relative to the unfiltered data. The effective overall resolution was calculated to be ∼55nm by Fourier ring correlation using NanoJ-SQUIRREL. Localized spot coordinates were reformatted and exported as .csv files for statistical analysis in the R software package SPATSTAT^62^. Post-filtered images typically contained 10^4^ – 10^5^ localizations/channel, and 5μm x 5 μm ROIs (in SLB-formed synaptic contacts) contained ∼ 5 x 10^3^ – 10^4^ localizations/channel. The *Lest* SPATSTAT core function was used to calculate autocorrelation of spots in an individual channel using Ripley’s L-function plotted as the normalized H function L(r)-r, as a function of r, where L is the L-function (variance-stabilized Ripley’s K-function derived by Besag’s transformation) and r is the correlation radius. This yields a value of 0 for a random Poisson distribution. Similarly, the *Lcross* core function was used to analyze cross-correlation of spots between two imaging channels, and plotted as the normalized H function. Negative values indicate dispersion (segregation) and positive values indicate coclustering. Images were rendered as Gaussian filtered images (using the Gaussian blur function in ImageJ – in which ‘intensity’ approximates point density) or as 2D point coordinates at higher digital magnification.

### Transmission electron microscopy

Sample processing of T cell/CHO cell conjugates for electron microscopy has been described in detail elsewhere. Briefly, conjugates of G8 T cell hybridomas and CHO cells expressing either T22 or T22-CD45 were prepared as for fluorescence microscopy. Culture medium was replaced with a mixture of 2.5% glutaraldehyde and 2% formaldehyde in 100mM cacodylate buffer (pH 7.4) with 2 mM Ca^2+^, post-fixed in buffered 1% osmium tetroxide and *en bloc* stained with 0.5% aqueous uranyl acetate. Ultrathin sections (∼60 nm thick) were cut and double-stained with uranyl acetate and lead citrate. Subsequent sample processing and image acquisition was performed as previously described. Intermembrane distance was measured in regions of the interface (at 200% digital magnification) where apposed membranes were aligned (parallel), and at which both membranes exhibited a trilaminar ‘tram-line’ appearance, indicating that they were orthogonal to the imaging plane.

### Statistical analysis

Unpaired T-test and One-way ANOVA (with correction for multiple comparisons) were performed to compare multiple sample means using GraphPad PRISM9 software. Pearson’s cross-correlation coefficient (PCC) is used to analyze colocalization in confocal *en face* synapse reconstructions, in which higher values indicate greater intensity colocalization between corresponding pixels from two images. For *d*STORM auto– and cross-correlation analysis averaged Ripley’s H function is plotted ± sem (see *dSTORM image processing*).

## Figures and Legends

**Extended Data Figure 1.**
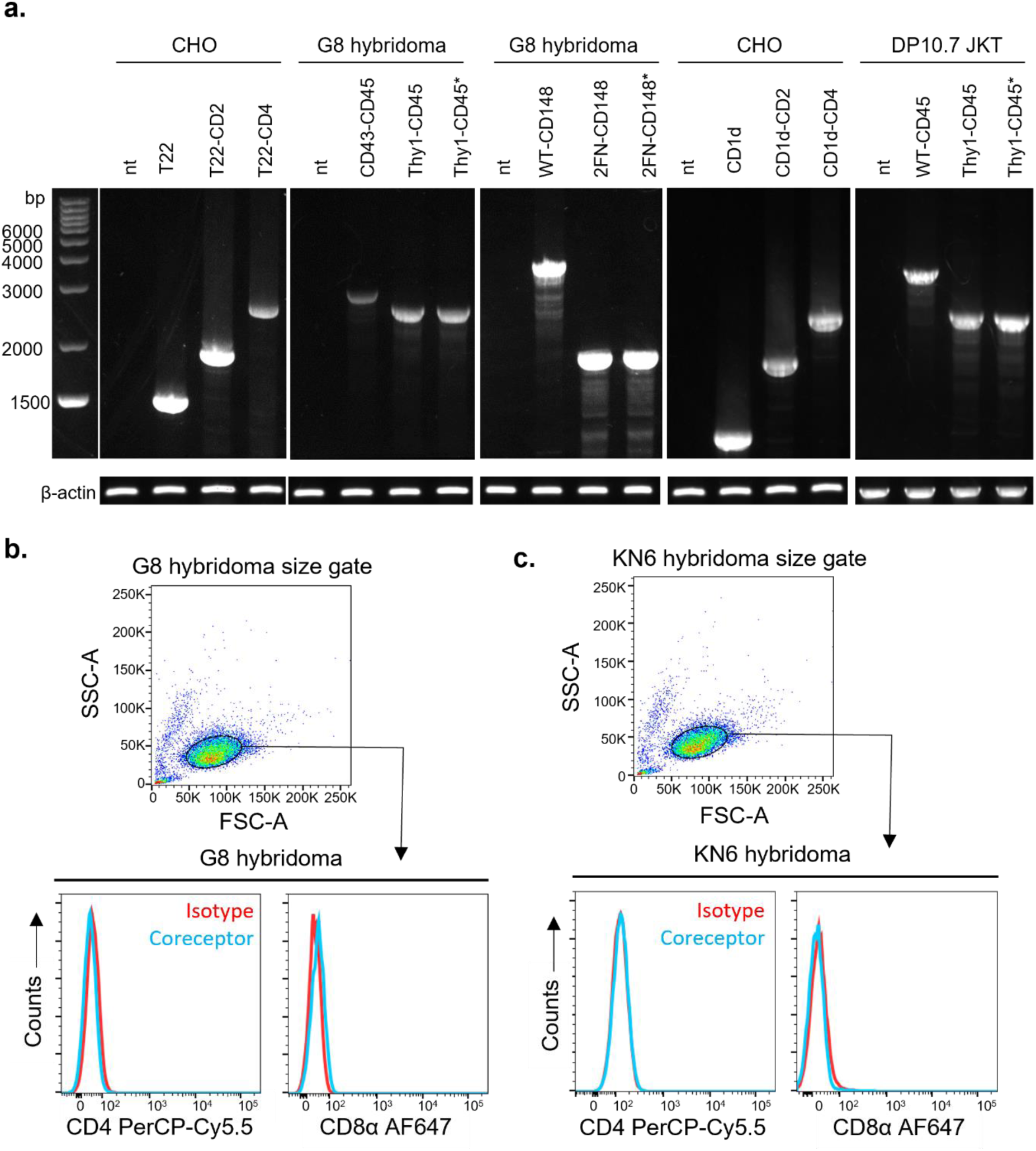
Characterization of CHO cell transductants and G8 T cell hybridomas. **a**. RT-PCR of total RNA extracted from the indicated cell lines stably transduced with the indicated constructs. RT-PCR reactions were performed using the primers listed in Extended Data Table 2. Histogram plots of surface expression of co-receptors CD4 and CD8 (cyan lines) in **b.** G8, and **c.** KN6, γδ T cell hybridomas, as measured by flow-cytometry. Red lines represent isotype control.

**Extended Data Figure 2.**
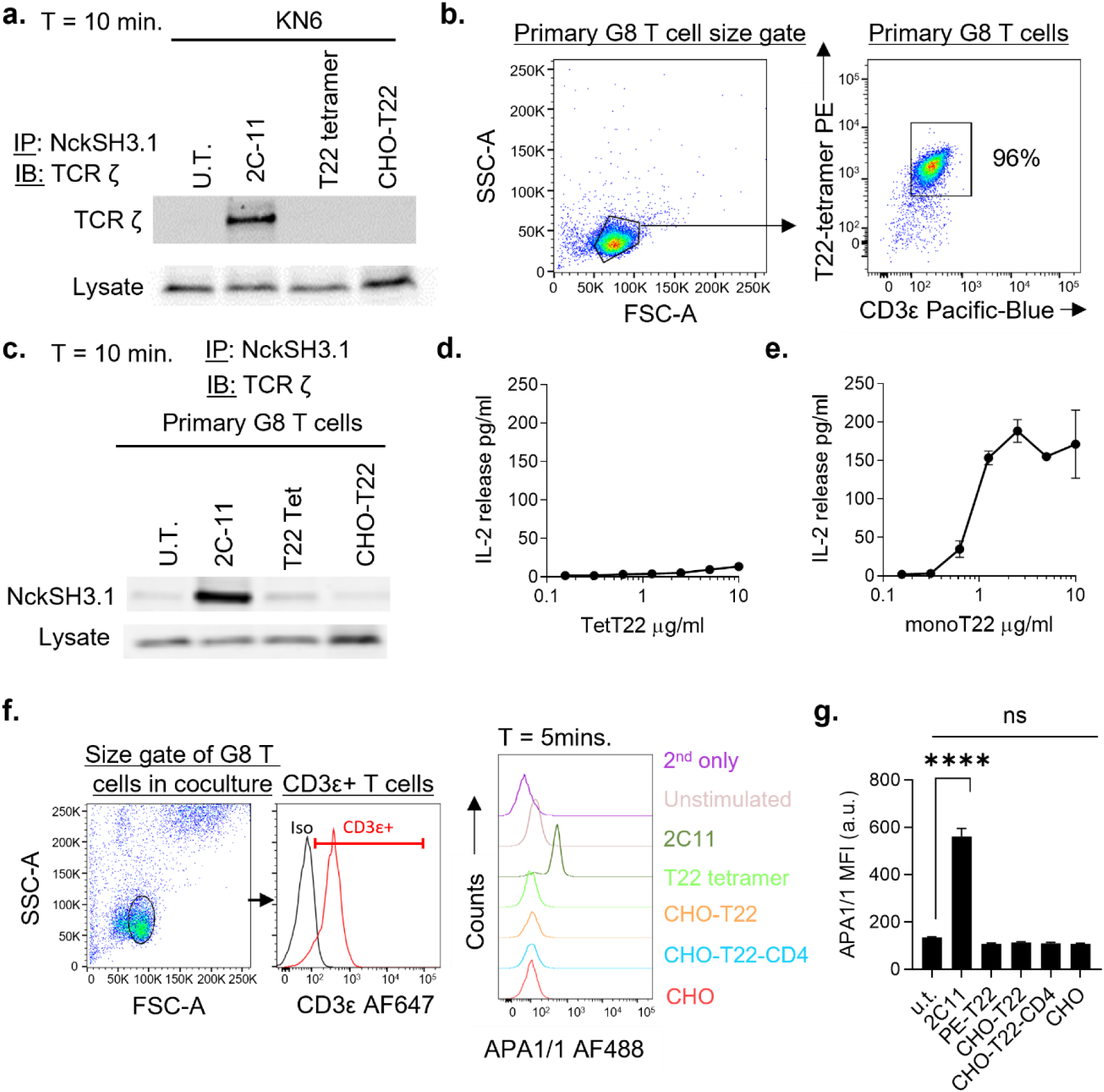
γδ TCR triggering does not require conformational changes typical of αβ TCRs. Detection of TCR conformational change by Nck SH3.1 pulldown assay and immunoblotting in **a.** KN6 γδ T cell hybridomas, or **c.** primary G8 γδ T cells, purified from G8 TCR transgenic mice (**b.**). Cells were incubated for 10 minutes at 37°C with 2C11 mAb, T22 tetramer, or T22-expressing CHO cells, prior to cell lysis and pulldown with biotinylated bead immobilised Nck SH3.1 peptide. Immunoprecipitated TCR in a binding-induced ‘open’ conformation capable of binding to Nck peptides was detected by SDS-PAGE and TCRζ immunoblot. Blots are representative of 2 (a.) and 3 (c.) independent experiments. **d.** IL-2 release by primary G8 T cells incubated with indicated concentrations of T22 tetramer for 24 hours. **e**. IL2-release by primary G8 T cells after 24hrs incubation in tissue culture wells pre-adsorbed with T22 monomers by overnight incubation at the indicated concentrations. **f.** Gating and histograms plots of primary G8 γδ T cells labelled with the conformation sensitive anti-CD3ε antibody APA1/1. Primary G8 T cells were stimulated as indicated, prior to fixation, permeabilization, and labelling with APA1/1 and a fluorescently labelled secondary antibody. **g.** Quantitation of labelling with APA1/1 antibody in *c.*, as measured by flow-cytometry. Data represent mean fluorescence intensity (MFI) of labelled cells treated is indicated. a.u., arbitrary units.; results are representative of 2 independent experiments. ****, P < 0.0001; ns, P > 0.05, after correction for all pairwise comparisons.

**Extended Data Figure 3.**
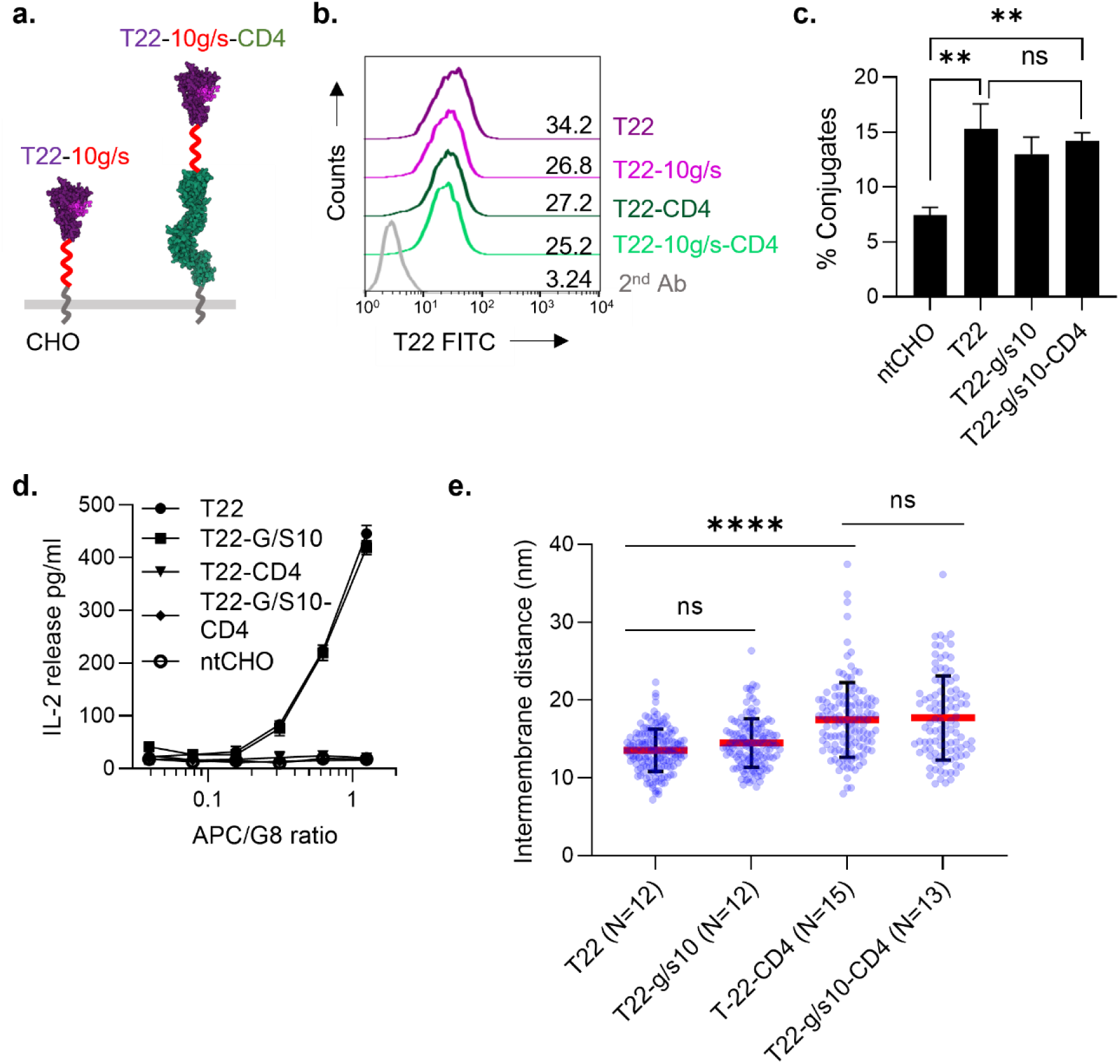
G8 T cell activation and intermembrane distance at G8 synapses is unaffected by increased T22 flexibility. **a**. Cartoon schematic of T22 and CD4-elongated T22 in which the 5 residue glycine/serine (g/s) linkers (grey) at the T22 C-terminus were replaced with 10 residue glycine/serine linkers (red) to increase flexibility of chimeric constructs. **b.** Histogram plots of CHO cell surface expression of the indicated T22 constructs as measured by flow-cytometry. Constructs were stably expressed in CHOs cells, and sorted for comparable expression. **c.** Cell-cell adhesion as measured by conjugate formation between G8 T cell hybridomas and CHO cells expressing indicated T22 constructs. Results are representative of 3 independent experiments. **d.** Activation of G8 γδ T cell hybridomas in response to original and flexible forms of T22, as measured by IL-2 release. 10^4^ G8 γδ T cell hybridomas were cocultured with increasing numbers of T22-expressing CHO cells for 24 hrs (expressed as APC/G8 cell ratio), and IL-2 release in culture supernatants measured by ELISA. Results are representative of 3 independent experiments. **e.** Intermembrane distances measured at synaptic contacts between G8 γδ T cell hybridomas and CHO cells expressing original and flexible versions of T22 and CD4-elongated T22. Data points are individual intermembrane distance measurements in each group. The number of conjugates sampled (N) is indicted for each group.

**Extended Data Figure 4.**
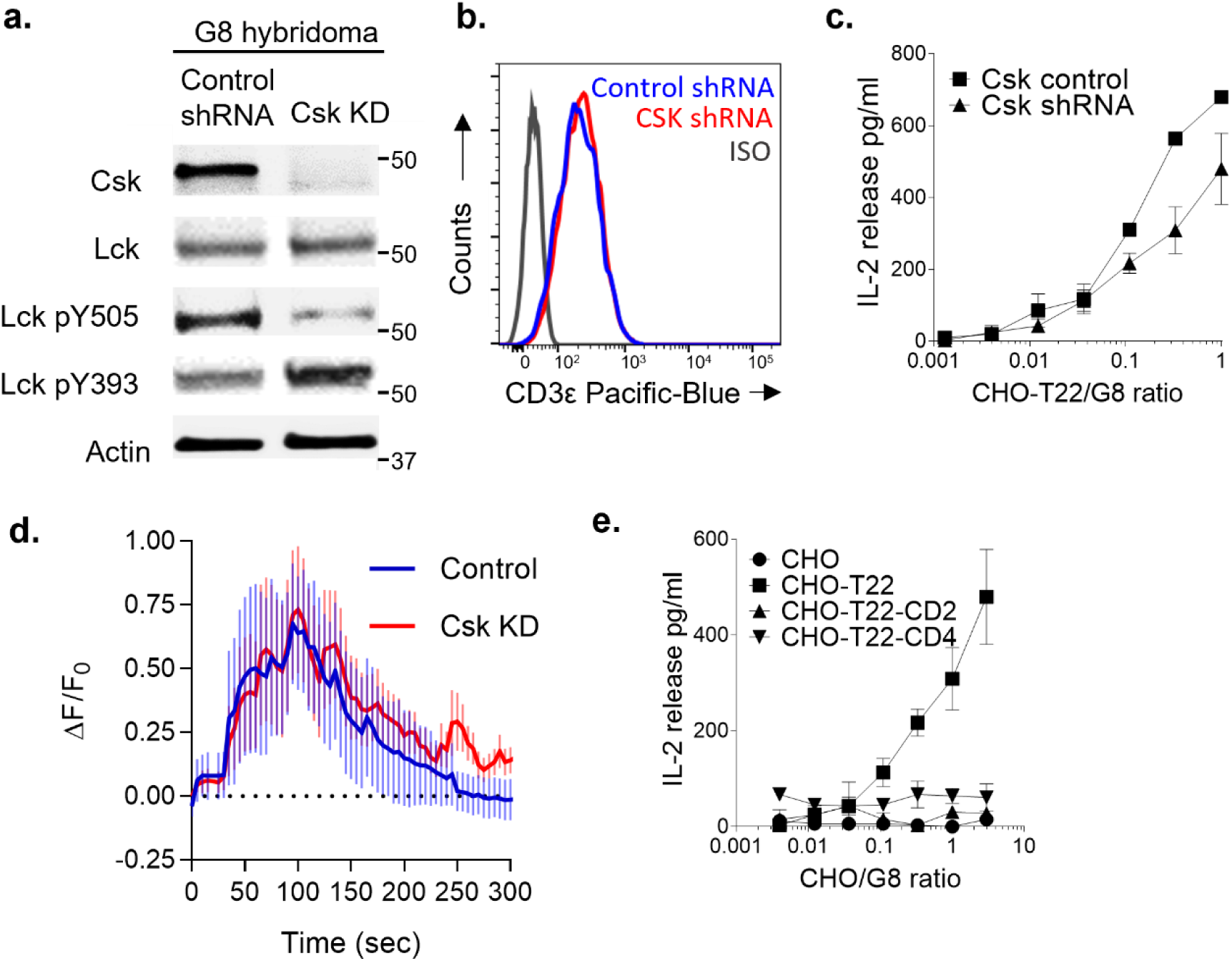
Suppression of Csk expression increases active Lck levels but fails to rescue TCR triggering in response to elongated T22. **a**. Immunoblots of resting G8 T cell hybridoma lysates that were stably transduced with Csk-targeting or non-targeting control shRNA. Cell lysates were analyzed using antibodies for Csk, and antibodies that specifically detect the tyrosine phosphorylated positive (Y393) and negative (Y505) regulatory sites of Lck. A representative blot from 4 independent experiments is shown. **b.** Cell-surface TCR expression levels of G8 hybridomas transduced with the indicated shRNA expression vectors shRNA. Cells were labeled with an anti-CD3ε antibody and analyzed by flow-cytometry. **c.** Activation of G8 T cell hybridomas stably transduced with Csk-targeting or control shRNA, as measured by IL-2 release after 24hrs co-culture with T22-expressing CHO cells. Results are representative of 3 independent experiments. **d.** Analysis of Ca^2+^ signaling by live G8 T cell hybridomas transduced with Csk targeting (Csk KD) or non-targeting (Control) shRNA following contact with planar lipid bilayers containing 100 molecules/μm^2^ T22 and 350 molecules/μm^2^ ICAM-1. T cells were loaded with Fluo-4 and Ca^2+^ flux imaged by epifluorescence microscopy following contact with supported lipid bilayers (T=0). Plots represent (mean fluorescence – fluorescence at T=0)/(fluorescence at T=0) ± SEM. Results are pooled from 2 independent experiments. **e.** IL-2 release in response to Increasing numbers of CHO cells expressing T22 and elongated forms of T22 (expressed as CHO/G8 ratio) by 10^4^ G8 T cell hybridomas expressing Csk-targeting shRNA. Results are representative of 3 independent experiments.

**Extended Data Figure 5.**
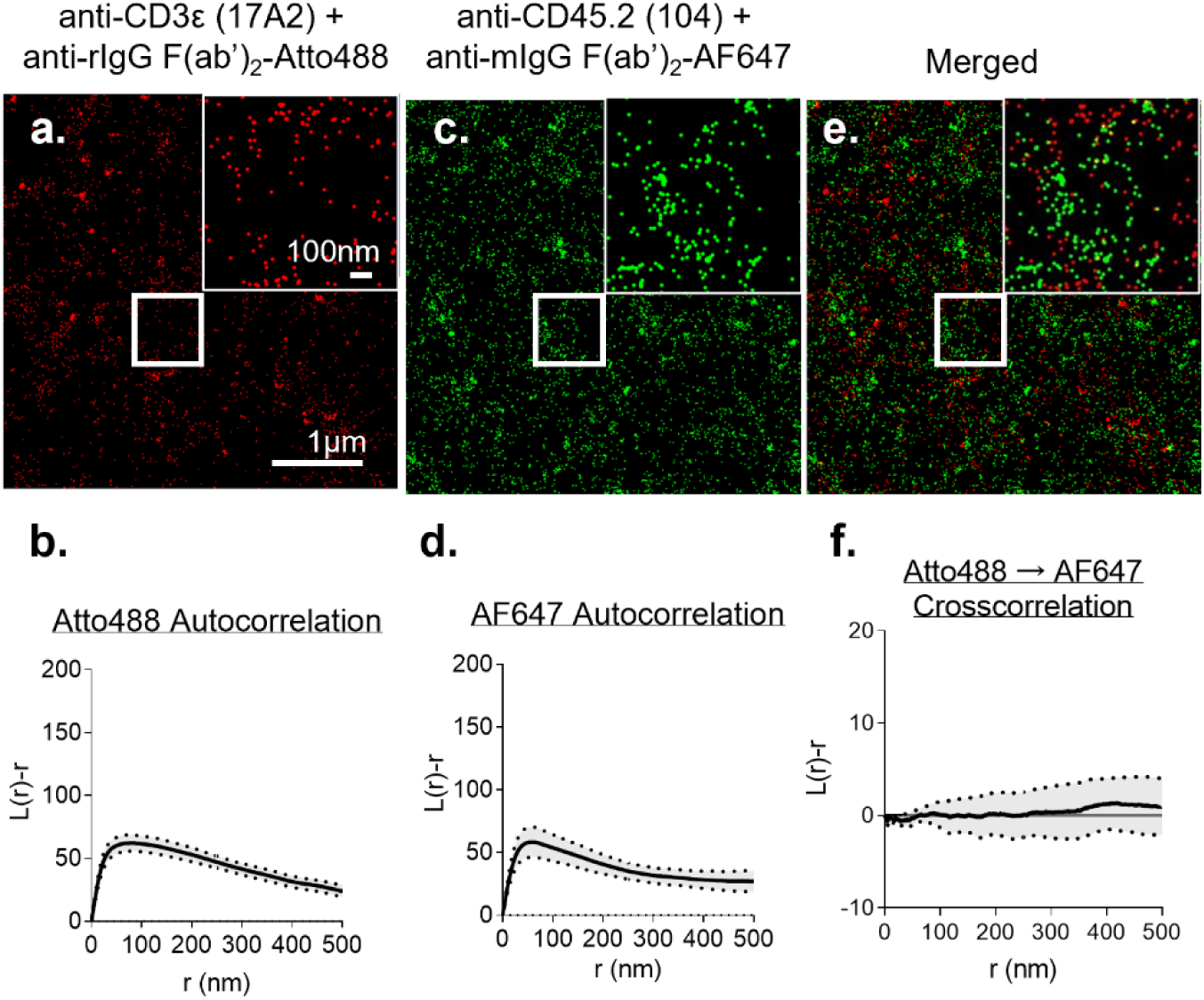
Characterization of antibodies used in *d*STORM imaging. **a**. *d*STORM localization image of rat anti-mCD3ε mAb 17A2 adsorbed onto a PLL-coated coverslip [indicated primary antibodies were adsorbed on PLL coverslip for 1hr at RT, fixed with 3% PFA for 45mins, quenched with 100mM glycine for 1hr, blocked with 5% BSA for 1hr, and stained with indicated 2nd antibody for 1hr (f/p ∼ 2)]. **b.** Ripley’s H-function autocorrelation of the Atto488 label. **c.** *d*STORM localization image of the anti-CD45.2 mAb 104 labelled with a polyclonal anti-mIgG (H+L chain) F(ab’)_2_-AF647 secondary antibody (f/p ∼2). **d.** Autocorrelation of the AF647 label. **e.** Merged image of Atto488 and AF647 localizations. **f.** Ripley’s H-function cross-correlation of Atto488 and AF647 labels. L(r)-r ∼0 for homgeneous random (Poisson) distribution, negative values represent dispersion (segregation) and positive values represent clustering. Shaded region within dotted lines demarcate 95% confidence intervals.

**Extended Data Figure 6.**
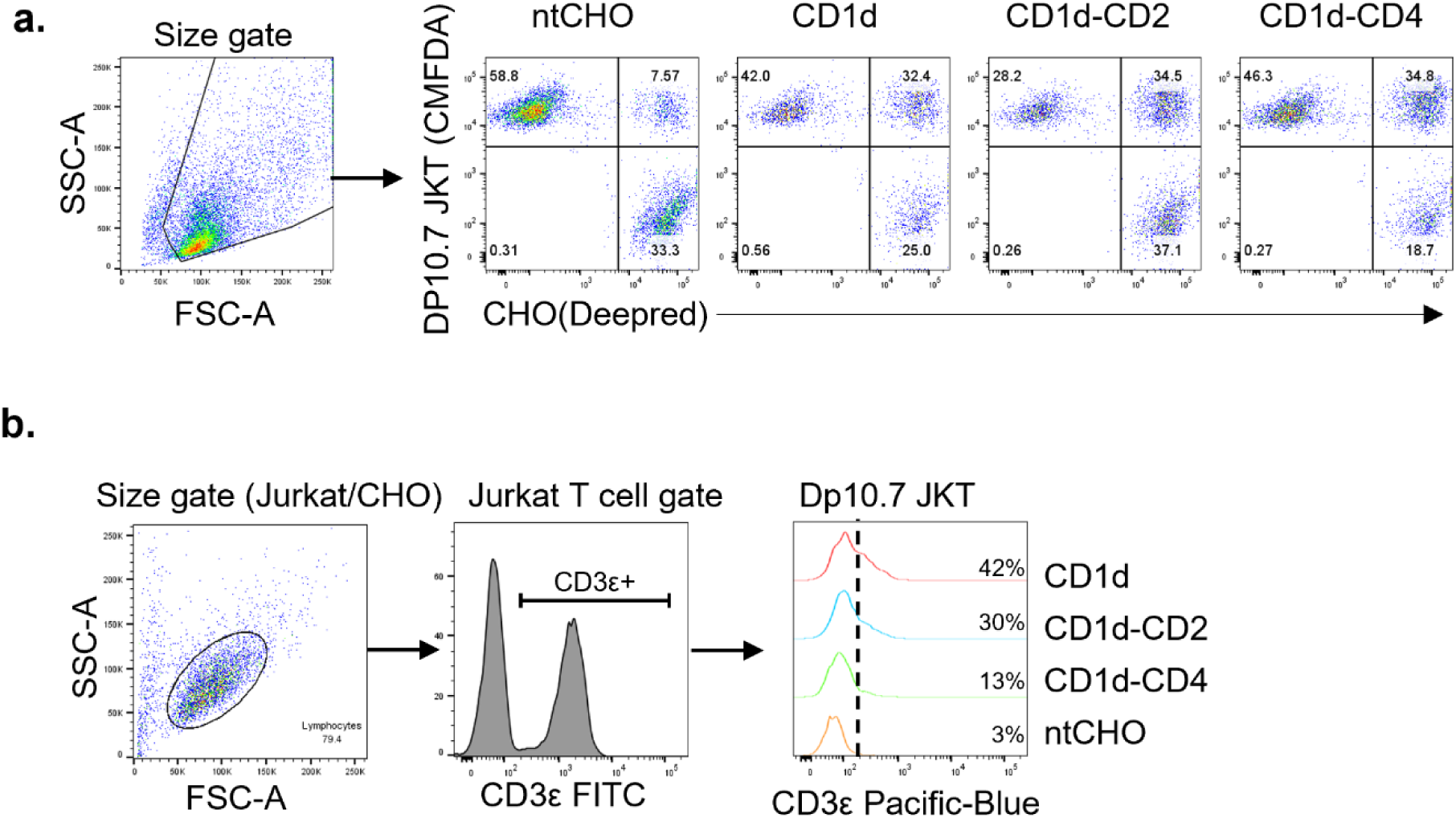
Flow-cytometry gating strategies. **a**. Adhesion assay in Figures 1 and 7, and **b.** CD69 upregulation in Figure 7, please also see *Methods*.

