## Supplementary Tables for "Ligand-induced segregation from large cell-surface phosphatases is a critical step in γδ TCR triggering"

**Supplementary Table 1. Quantitation of cell surface molecules**

| Cell Type | Construct expressed | mol/μm$^2$ | % of endogenous CD45/CD148 |
|---|---|---|---|
| G8 T cell hybridoma | None (Endogenous CD45) | 793.8 | 100 |
| | Thy-CD45 | 3.6 | 0.5 |
| | Thy1-CD45* | 2.9 | 0.4 |
| | CD43-CD45 | 16.3 | 2.1 |
| | None (Endogenous CD148) | 43.8 | 100 |
| | 2FN3-CD148 | 19.4 | 44.3 |
| | 2FN3-CD148* | 18.8 | 42.3 |
| | WT-CD148 | 3.2 | 7.3 |
| DP10.7 Jurkat T cell | None (Endogenous CD45) | 622 | 100 |
| | Thy-CD45 | 34.1 | 5.3 |
| | Thy1-CD45* | 72.2 | 11.2 |
| | WT-CD45 | 38.0 | 5.9 |

*Catalytically inactive mutant

**Supplementary Table 2. Oligonucleotide sequences**

| | | |
|---|---|---|
| T22 constructs | T22 | Forward: tcgggtggcggcggctctggttcacactcgcttaggtatttc |
| | | Reverse: gaggaagcttccagtcaaggtgacagtaaagactcg |
| | β-2M | Forward: gatatctcgagtcgcttcagtcgtcagcatgg |
| | | Reverse: ccagagccgccacccgagccgcctccgcccgaaccgcca cctcccatgtctcgatcccagtagac |
| | BamHI site insertion | Forward: taatggatcctgcatggaaagtggtgc |
| | | Reverse: taatggatcctcagtcggcagaacctg |
| | BamHI site removal | Single primer: gatgggaggatccgtggatttggattgttg |
| | CD45 STOP insertion | Forward: gagcaagggcgagtagctgttcaccgg |
| | | Reverse: ccggtgaacagctactcgcccttgctc |
| RT-PCR | T22-SCD | Forward: atggctcgctcggtgacc |
| | | Reverse: aggtgacagtaaagactcgcca |
| | Flag-mouseCD45 | Forward: gattacaaggatgacgacgataagccta |
| | | Reverse: cgtgaactctgggttggagct |
| | Flag-mouseCD148 | Forward: gattacaaggatgacgacgataagccta |
| | | Reverse: gatgtaaccattagtctttccaaacatgct |
| | CD1d-SCD | Forward: ggggggatcccggtactcgagggcttcaataag |
| | | Reverse: agtcgcggccgctcacaggacgccctgatagga |
| | Flag-humanCD45 | Forward: gggattataaggacgatgatgacaaaca |
| | | Reverse: tgaaccttgatttaaagctggacttg |
| | Mouse β-actin | Forward: ctgtccctgtatgcctctg |
| | | Reverse: atgtcacgcacgatttcc |
| | Hamster β-actin | Forward: ctgtccctgtatgcctctg |
| | | Reverse: atgtcacgcacaatttcc |
| | Human β-actin | Forward: gagcacagagcctcgccttt |
| | | Reverse: ccaggaaggaaggctggaag |
